# Neural representations in visual and parietal cortex differentiate between imagined, perceived, and illusory experiences

**DOI:** 10.1101/2023.03.31.535014

**Authors:** Siyi Li, Xuemei Zeng, Zhujun Shao, Qing Yu

## Abstract

Humans constantly receive massive amounts of information, both perceived from the external environment and imagined from the internal world. To function properly, the brain needs to correctly identify the origin of information being processed. Recent work has suggested common neural substrates for perception and imagery. However, it has remained unclear how the brain differentiates between external and internal experiences with shared neural codes. Here we tested this question by systematically investigating the neural processes underlying both the generation and maintenance of information from voluntary imagery, veridical perception, and illusion. The inclusion of illusion allowed us to differentiate between objective and subjective internality: while illusion has an objectively internal origin and can be viewed as involuntary imagery, it is also subjectively perceived as having an external origin like perception. Combining fMRI, eye-tracking, multivariate decoding and encoding approaches, we observed superior orientation representations in parietal cortex during imagery compared to perception, and conversely in early visual cortex. This imagery dominance gradually developed along a posterior-to-anterior cortical hierarchy from early visual to parietal cortex, emerged in the early epoch of imagery and sustained into the delay epoch, and persisted across varied imagined contents. Moreover, representational strength of illusion was more comparable to imagery in early visual cortex, but more comparable to perception in parietal cortex, suggesting content-specific representations in parietal cortex differentiate between subjectively internal and external experiences, as opposed to early visual cortex. These findings together support a domain-general engagement of parietal cortex in the generation and maintenance of internally-generated experience.

**Significance Statement:** How does the brain differentiate between imagined and perceived experiences? Combining fMRI, eye-tracking, multivariate decoding and encoding approaches, the current study revealed enhanced stimulus-specific representations in visual imagery originating from IPS, supporting the subjective experience of imagery. This neural principle was further validated by evidence from visual illusion, wherein illusion resembled perception and imagery at different levels of cortical hierarchy. Our findings provide direct evidence for the critical role of parietal cortex as a domain-general source region for the generation and maintenance of content-specific imagery, and offer new insights into the neural mechanisms underlying the differentiation between subjectively internal and external experiences.

## Introduction

In complex environments, humans are overwhelmed with information from external and internal origins. To function properly, the brain needs to differentiate between what is perceived externally and what is imagined internally. Extensive evidence has suggested a large overlap in neural processing between imagery and perception, in terms of common univariate BOLD activations (Ishai et al., 2000; Ganis et al., 2004), and of shared neural representations between perceived and imagined contents as revealed by multivariate decoding approaches in visual cortex (Stokes et al., 2009; Reddy et al., 2010; Albers et al., 2013; Ragni et al., 2020). Yet, it is less understood why imagery and perception feel so different, despite these similarities (Koenig-Robert and Pearson, 2021).

Previous work has demonstrated a reduced level of neural activation (Ishai et al., 2000; Ganis et al., 2004; Keogh et al., 2020), as well as weaker stimulus-specific representations (Lee et al., 2013; Dijkstra et al., 2018), during imagery in early visual cortex (EVC) as compared to perception, given that perception is driven by sensory signals reaching first at EVC, whereas imagery does not receive any direct external stimulations (Iamshchinina et al., 2021; Yu and Postle, 2021). This account could possibly explain why perception evokes a stronger sensory experience than imagery, but does not explain how imagery contents are generated in the absence of external stimulation, and why imagery feels internal. Although previous work has demonstrated top-down feedback modulation from frontoparietal cortex during imagery (Mechelli et al., 2004), most of the observations were based on univariate activations, and do not speak to whether these higher-order brain regions carry content-specific neural representations that discriminate between imagery and perception. Here, we tested the idea that neural signals contributing to the subjective internality of imagery arise from higher-order brain regions that serve as the source of content-specific imagery representations. As discussed earlier, external experience of perception is supported by stronger neural representations in EVC; with this logic, we propose that content-specific imagined representations should exceed perceptual representations in (some) higher-order regions, resulting in the internal experience of imagery. Moreover, given such a neural principle exists, it should be generalizable to conditions which are neither completely internal nor external, such as illusion (Bergmann et al., 2019). Illusion is another type of sensory experience that is dissociated from direct sensory inputs, and can therefore be conceptualized as involuntary imagery (Pearson, 2019). On the other hand, similar to veridical perception, illusion creates the subjective feeling that information is stimulated externally. In other words, illusion and perception can be considered as subjectively external experiences, while imagery is subjectively internal. We therefore predict that illusion should share neural similarities with perception and imagery at different levels of cortical processing: it should behave more similarly as imagery at the early stage of cortical processing, and should resemble perception to a larger extent at the later stages.

Here we propose that parietal cortex remains a good candidate for the source region of content-specific imagery, for several reasons: first, although earlier research suggested variable representations of imagined contents in parietal cortex, recent advances in visual working memory research have suggested robust stimulus-specific information in working memory in parietal cortex (Sprague and Serences, 2013; Ester et al., 2015; Yu and Shim, 2017). Given that working memory and imagery are tightly linked (Albers et al., 2013), it would be reasonable to expect robust stimulus-specific representations in parietal cortex during imagery. Second, parietal cortex encodes various types of information during visual imagery and working memory, including spatial (Sprague and Serences, 2013), feature (Yu and Shim, 2017), object (Ragni et al., 2020) and pictorial (Breedlove et al., 2020) information. Lastly, parietal cortex also encodes information in various illusory contexts (Liu et al., 2019; Arsenovic et al., 2022). Despite all these, surprisingly, none of the previous studies, to our knowledge, have directly tested whether representations of imagery exceed those of perception and illusion in any brain regions, including parietal cortex.

## Materials and Methods

### Participants

A total of 52 volunteers participated in the study. All were recruited from the Chinese Academy of Science Shanghai Branch community, and were naïve to the purpose of the study. Seventeen volunteers (6 males, mean age = 23.4 ± 1.5 years) participated in Experiment 1. Twenty-one volunteers (9 males, mean age = 24.9 ± 1.7 years) participated in the main experiment of Experiment 2. Five volunteers (2 males, mean age = 24.0 ± 1.4 years) participated in the retrocue version of Experiment 2. Seventeen volunteers (3 males, mean age = 23.8 ± 1.8 years) participated in the eye-tracking version of Experiment 2. Several volunteers participated in more than one experiment: one volunteer participated in both main experiments of Experiments 1 and 2. All volunteers in the retrocue version also participated in the main experiment of Experiment 2. One volunteer participated in both Experiment 1 and eye-tracking experiment, and one volunteer participated in both main experiment of Experiment 2 and eye-tracking experiment. For main experiment of Experiment 2, three participants were excluded due to excessive head motion or technical problems during fMRI scanning, leaving 18 participants in the final sample (8 males, mean age = 24.8 ± 1.8 years). For eye-tracking experiment, four participants were excluded due to excessive eye blinks/head motion or technical problems during recording, resulting a final sample size of 13 (2 males, mean age = 23.7 ± 1.6 years). We did not determine sample size of each experiment a priori, but the sample size used in the current study was comparable to those in studies with a similar approach (Ester et al., 2015; Liu et al., 2019; Yu and Postle, 2021).

All participants had normal or corrected-to-normal vision and reported no neurological or psychiatric disease. All participants were provided written, informed consent approved by the ethics committee of the Center for Excellence in Brain Science and Intelligence Technology, Chinese Academy of Sciences, and were monetarily compensated for their participation. Participants filled out a revised version of the vividness of visual imagery questionnaire (VVIQ) (Marks, 1973) as an evaluation of their general imagery ability (1-5 points of rating: 1 – “no image at all”; 5 – “as vivid as normal vision”), and those who reached an average VVIQ score of above 2 (Kay et al., 2022) moved onto the main experiment. All of our participants met this criterion.

### Stimuli and Procedure

#### Overview

The current study investigated neural processes underlying the generation and maintenance of stimulus-specific information during voluntary imagery, veridical perception, and illusion. Typical imagery tasks often involve a delay period during which imagery contents emerge and persist, whereas perception does not. Hence, instead of simply contrasting memory delay of imagery and sample encoding of perception that pertain to two temporally-distinct processes, we embedded a delay period into each condition, which would allow comparisons of neural codes between distinct processes. In all three conditions, the trial started with the presentation of a perceptual or illusory stimulus, or an imagery cue, followed by a common delay period before a response was made. We referred these three conditions as perception-based, imagery-based, and illusion-based delayed-recall, respectively (perception, imagery and illusion in short, respectively). We used two distinct sets of stimuli to test our hypothesis: moving Gabors along oriented paths in Experiment 1 and static line orientations in Experiment 2, and the way to cue imagery also differed between experiments. Below we presented details for the generation of stimuli and experimental procedure in both experiments.

For fMRI, all stimuli were generated using MATLAB R2012b (The MathWorks) and Psychtoolbox-3. Stimuli were presented on a SINORAD monitor screen (37.5×30 cm, 1280×1024 pixels at 60 Hz) at the back of the scanner bore and projected onto a mirror mounted on the head coil, with a viewing distance of 90.5 cm. Participants lay supinely in the scanner and completed the task using two SINORAD two-key button boxes. Eye-tracking was performed using MATLAB R2018a, a viewing distance of 70 cm, and screen resolution of 1680×1050.

#### Experiment 1

In Experiment 1, stimuli were Gabor patterns (a full contrast sinusoidal grating weighted by a Gaussian function) with a spatial frequency of 0.94 circles per degree and a size of 1.6° of visual angle in diameter, presented against a uniform gray background (RGB values: [128, 128, 128]). On perception trials, the Gabor pattern moved back and forth along a tilted (leftward or rightward relative to the vertical line) linear path with its internal grating texture remaining static. The motion direction of the Gabor reversed every 1 sec and the path orientation was determined by an illusion size measurement task (see below), separately for each individual prior to scanning. The speed in the vertical direction was 5°/sec whereas the speed in the horizontal direction and the path length could vary across subjects, depending on the orientation of the path. On imagery trials, the stimulus was replaced by a symbolic cue centered at the fixation point. The letter ‘L’ and ‘R’ were used to, respectively, prompt subjects to form imagery of a Gabor moving back and forth along a leftward and rightward pathway (relative to the vertical line) without actually viewing the Gabor pattern (for two of the participants, a light gray and a dark gray circle were used as cues instead of letters, and were presented peripherally at the center of the motion path). On illusion trials, the Gabor pattern moved back and forth vertically along a linear path with a speed of 5°/sec (external motion), and, simultaneously, its internal texture drifted in a horizontal-left or horizontal-right direction with a temporal frequency of 4 cycles/sec (internal motion), making a double-drift stimulus (Liu et al., 2019). By presenting the stimulus in the participant’s peripheral visual field, the combination of the external motion and the orthogonal internal motion would tilt the perceived motion path either to the left or right relative to the physical motion path (i.e., inducing a visual illusion of the path orientation). The length of the motion trajectory was 5°, and the direction of the internal motion reversed at the two endpoints, in synchrony with path reversals, every 1 sec. For perception and illusion conditions, the midpoint of the motion trajectory was located at 5° horizontal to the right of the screen center, and the initial spatial phase of the Gabor pattern was randomly generated for each trial. Moreover, a fixation point (0.3° diameter) in black was displayed at 3° horizontal to the left of the screen center, making the stimulus appear in a more peripheral visual field (8° of eccentricity) for participants (Figure 1A).

**Figure 1.**
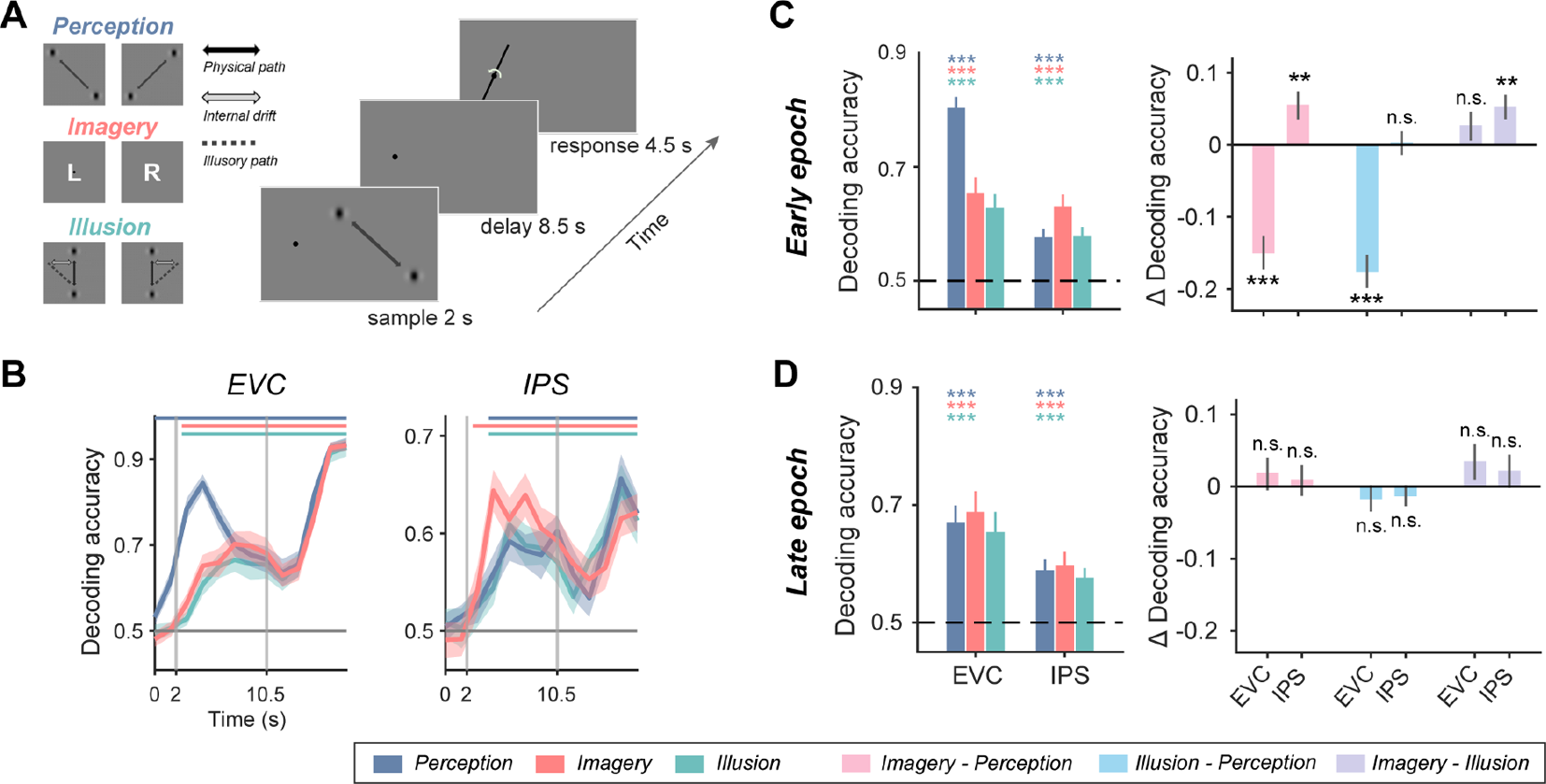
Task design and orientation representations in Experiment 1. A. In Experiment 1, participants (n = 17) performed a delayed-recall task, during which they precisely memorized a physical, imagined, or illusory orientation of the motion path of a Gabor pattern over a prolonged delay. Each trial began with the presentation of one sample stimulus (leftward or rightward) from one of the conditions in the visual periphery. After a delay, participants rotated the needle presented at fixation to match the orientation of the remembered path as precisely as possible. B. Time course of decoding accuracy in perception (blue), imagery (red), and illusion (green) conditions, in EVC and IPS of Experiment 1. ROIs were contralateral to the side of stimulus presentation. Colored lines on top denote significant time points of the corresponding condition. Vertical gray lines denote onset of delay (at 2 s) and of probe (at 10.5 s). Horizontal lines denote chance level of 0.5. Shaded areas denote error bars (±1 SEM). C. Left panel: average decoding accuracy in perception, imagery, and illusion conditions, in early epoch of EVC and IPS of Experiment 1. Horizontal dashed lines denote chance level of 0.5. Error bars denote ±1 SEM. Colored asterisks on top denote significance of the corresponding condition. Right panel: Differences in decoding accuracy, between imagery and perception (pink), between illusion and perception (light blue), and between imagery and illusion (purple), in early epoch of EVC and IPS of Experiment 1. Error bars denote ±1 SEM. Asterisks denote significance of pairwise comparisons between conditions. n.s.: not significant, *: *p* < 0.05, **: *p* < 0.01, ***: *p* < 0.001. D. Same as C, but with results from late epoch.

Prior to the main experiment, participants were measured on the size of their double-drift illusion inside the scanner. Stimuli parameters were identical to those used in the main task. Participants were instructed to keep their gaze at the fixation point throughout all tasks, unless specified. Each trial began with a 1-sec fixation period. Following that, a Gabor pattern appeared in the periphery and moved back and forth along a vertical path for 2 sec, with its internal texture drifting in an orthogonal direction (double-drift left-tilted and double-drift right-tilted trials) or remaining static (no-drift trials). After that, a response needle (0.05° in width, 5° in length) centered at the fixation point was presented. Participants used four buttons to rotate the needle until it matched their perceived angle of the motion trajectory of the Gabor pattern as precisely as possible. The initial orientation of the needle was randomly chosen on every trial. Participants made clockwise or counterclockwise rotations in steps of 1° by pressing one of two buttons (labelled with ‘1’, ‘2’) on the right button-box with their thumb, and could quickly rotate the bar by 90° via pressing the ‘3’ button on the left button-box. Participants pressed the ‘enter’ button (labelled with ‘4’) on the left button-box to indicate they were satisfied with their response and to start the next trial. In each run, three types of stimuli (left-drift, right-drift, and no-drift of the Gabor internal texture) were randomized across trials and each type was presented on one-third of 30 trials. Twelve participants completed 2 runs of the measurement task and five participants completed 3 runs, lasting 9-14 minutes in total. To ensure accurate responses, only data from the last run was used to calculate the orientation of the physical path of the perception stimulus in the main task. Participants practiced 2-3 runs of the measurement task during a behavioral practice session prior to the fMRI session.

On every trial of the delayed-recall task, the sample stimulus (a symbolic cue or a moving Gabor pattern) was displayed for 2 sec, followed by an 8.5-sec delay. During perception and illusion trials, participants viewed the moving Gabor pattern displayed in their right peripheral visual field and remembered it over the delay. During imagery trials, participants were prompted by the cue to actively imagine a Gabor moving back and forth along the corresponding (leftward or rightward) trajectory during delay. Following the delay, participants rotated a needle (0.05° in width, 5° in length) centered at the fixation point to make it match the orientation of the trajectory of the memorized or imagined moving Gabor pattern. Participants had 4.5 sec to respond using the same four buttons as in the measurement task. Each trial ended with an inter-trial interval (ITI) of 4.5, 6 or 7.5 sec. In each run, participants performed 18 trials (6 min and 25.5 sec per run), and the conditions (imagery, perception and illusion) and stimulus directions (i.e., leftward or rightward trajectory) of trials were balanced and randomly ordered. Overall, sixteen participants completed 14 runs (i.e., 252 trials in total) within a single fMRI experiment session lasting around 120 minutes, except one subject (S002) who completed 12 runs due to a technical problem with the scanner. Before the fMRI session, participants practiced 1-2 runs of the delayed-recall task outside the scanner to make sure that they correctly understood the task instruction and were comfortable with the button responses.

#### Experiment 2

In Experiment 2, sample stimuli were oriented bars randomly chosen from a continuous orientation space from 1° to 180° in steps of 1°. On perception trials, the stimulus was made up of two distant discs (0.8° in diameter) and a bar (0.6° in width) connecting them; whereas on imagery trials, the stimulus consisted of two distant discs only. Thus, participants memorized a physically-present oriented bar on perception trials but need to imagine the missing bar between two discs on imagery trials. Inspired by the classic Kanizsa triangle illusion, the stimulus on illusion trials consisted of two distant discs with opposite openings facing each other (Figure 4A), which caused a subjective visual experience of a completed illusory contour defining an oriented bar that occluded discs at both ends (i.e., seeing an outline that was nonexistent in fact). All stimuli were in black and presented centrally on a uniform grey background screen (RGB values: [128, 128, 128]) with a radius of 4°. To prevent participants from memorizing the location of the end discs instead of the bar orientation, a jitter of ± 0.4° was applied to the radius on every trial.

The trial time course of the delayed-recall task in Experiment 2 was similar to that in Experiment 1: on each trial, a sample stimulus with a randomly-selected orientation was displayed for 1 sec without a fixation point. A 9.5-sec delay then followed. During perception and illusion trials, participants memorized the perceived oriented bar throughout the delay, while during imagery trials, participants were required to internally generate the mental image of a bar connecting two end discs and keep it in their mind’s eye. Following that, a probe dial (4° radius) appeared centrally with its needle in a random initial orientation. Participants rotated the needle to match the maintained orientation in a 4.5-sec response window. The usage of response buttons was identical to that in Experiment 1. ITI varied in 6, 7.5 or 9 sec. At the end of each run, feedback regarding participants’ overall performance was provided. Each run consisted of 18 trials (6 of each condition) in a randomized order, lasting 6 min and 52.5 sec. Participants completed 20 runs of the delayed-recall task across two fMRI scanning sessions, and the orientations of sample stimuli evenly covered the entire orientation space across 10 consecutive runs within every session. In total, seventeen participants completed 360 trials (i.e., 120 trials for each condition), and one participant (S012) completed 342 trials (i.e., 19 runs). Moreover, 5 runs for S006 were not brought into subsequent analyses due to a failure in alignment, so were 10 runs for S021 due to a technical problem with the scanner. Prior to the fMRI sessions, participants underwent a practice session. Practice session ended when the mean absolute response error of the last practice run fell below 15°.

In addition to the delayed-recall task, Experiment 2 also included an independent perception-mapping task. The stimuli of this task were identical to the sample stimuli of perception trials in the main task. On every trial, a stimulus (an oriented bar with two end discs) flickering at 1.8 Hz was presented for 4.5 sec, followed by an ITI of 6, 7.5 or 9 sec. Each run consisted of 30 trials and lasted 4 min and 37.5 sec. Throughout the run, participants performed a detection task to maintain their attention on the fixation point, during which they needed to press the ‘1’ button as quickly as possible whenever the fixation point turned green. Data of six runs of the perception-mapping task were acquired across two fMRI scanning sessions for seventeen participants, and stimuli of 180 trials evenly covered 1° to 180°; and data of five runs were obtained for one participant (S012). For one participant (S021), data of three runs in the first session were not included in subsequent analyses due to a technical problem with the scanner. Participants also practiced 2-3 runs to reach an accuracy of above 70% prior to scanning.

In the retrocue version of Experiment 2, the trial time course was similar to that in the main experiment of Experiment 2, except that two sample stimuli were presented, followed by a retrocue, before the onset of the delay. Specifically, each sample was presented for 0.8 sec with an inter-stimulus-interval of 0.4 sec. After 0.4 sec following the offset of samples, a retrocue (a digit of “1” or “2”) was presented for 0.6 sec. Participants needed to maintain the cued sample during the delay. The response window was 4.5 sec, and ITI varied in 4.5, 6, and 7.5 sec. Each run consisted of 18 trials, and participants completed 20 runs across two scanning sessions.

In the eye-tracking version of Experiment 2, the trial time course and stimuli were also similar to those in the main experiment of Experiment 2, except that sample and delay periods were shortened to 0.4 and 2.4 sec, respectively, and the radius of stimuli was changed to 2.5°. Trials from different conditions were conducted in separate runs in a randomized order instead of interleaved in one run. Each run consisted of 60 trials, and participants performed six runs in total. Participants’ eye gaze positions were recorded using a Desktop mount Eyelink 1000 Plus eye tracker (SR Research) at 1000 Hz.

### Behavioral Analyses

For illusion size measurement in Experiment 1, the perceived orientation of each trial was calculated as the absolute circular distance between the response orientation and the vertical line, then the difference between the mean of perceived orientation on double-drift and that on no-drift trials was calculated as the orientation of the physical path on perception trials in the main task (i.e., the anchor orientation). For the delayed recall task in Experiment 1, recall error was computed as the angular difference between the reported and anchor orientations. Moreover, the standard deviation (SD) of the reported orientations (relative to the vertical line) of each condition were computed to indicate recall variability. Differences between conditions were characterized using one-way repeated measures ANOVAs. The behavioral data of three participants (S001, S002 and S003) were not recorded due to a coding error, so behavioral results for Experiment 1 reported here were based on data of 14 participants. Similar analyses were performed for recall error and RT in Experiment 2, in which recall error was computed as the absolute circular distance between the response orientation and sample orientations.

### Data Acquisition

MRI scanning was performed at the Center for Excellence in Brain Science and Intelligence Technology (CEBSIT), Functional Brain Imaging Platform (FBIP) on a Siemens 3T Tim Trio MRI scanner with a 32-channel head coil. High-resolution T1-weighted anatomical images were acquired using a magnetization-prepared rapid gradient-echo (MPRAGE) sequence (2300 ms time of repetition [TR], 3 ms time of echo [TE], 9° flip angle [FA], 256 × 256 matrix, 192 sequential sagittal slices, 1 mm^3^ isotropic voxel size). Whole-brain functional images were acquired using a Multiband 2D gradient-echo echo-planar (MB2D GE-EPI) sequence with a multiband factor of 2, 1500 ms TR, 30 ms TE, 60° FA within a 74 × 74 matrix (46 axial slices, 3 mm^3^ isotropic voxel size). For three participants (S001, S006, and first session of S005), functional images were acquired using the MB2D GE-EPI sequence with a FA of 90° and other parameters remained same as above.

### fMRI Preprocessing

Preprocessing of fMRI data was performed using AFNI (afni.nimh.nih.gov; (Cox, 1996)). The first five volumes of each functional run were removed. Following that, data from each scanning session were first registered to the last volume of the last run in the same session and then to the T1-weighted anatomical images obtained in the first session. After within-session (Experiment 1) or cross-session (Experiment 2) alignment, motion correction and detrending (linear, quadratic, cubic) were applied to the data. Voxel-wise signal timeseries were z-scored on a run-by-run basis for subsequent univariate and multivariate analyses.

### ROI Definition and Voxel Selection

We used the probabilistic atlas developed by Wang and colleagues (Wang et al., 2015) to identify the individual anatomical regions of interest (ROIs). Masks of the probabilistic atlas were first warped back to each participant’s structural images in their native space, including V1, V2, V3, hV4, V3AB, VO1, VO2, hMT, MST, IPS0-5, and FEF. After that, we extracted the unilateral masks of V1, V2 and V3 of each participant and merged them within (for Experiment 1) or between (for Experiment 2) hemispheres to create the anatomical early visual cortex (EVC) ROIs. Similarly, we merged masks of IPS0-5 create the anatomical intraparietal sulcus (IPS) ROIs, merged masks of hMT and MST to create the anatomical MT+ ROIs, and used mask of FEF to create the anatomical superior precentral sulcus (sPCS) ROIs.

To create the functionally-defined ROIs, we conducted voxel selection within each anatomical ROI based on task-related activations. A conventional mass-univariate general linear model (GLM) analysis was implemented in AFNI, where the GLM included six nuisance regressors regarding the participant’s head motion and three regressors of interest: sample, delay, and probe periods of the task modelled with boxcars (with a duration of 2 s, 8,5 s, and 4.5 s, respectively for Experiment 1, and a duration of 1 s, 9.5 s, and 4.5 s, respectively for Experiment 2) that were convolved with a canonical hemodynamic response function. The ‘sample – baseline’ contrast was used to define the functional EVC ROIs, and the ‘delay - baseline’ contrast was used to define the rest of the functional ROIs, by selecting 500 voxels that displayed the strongest responses on the specific contrast within the anatomical ROI. Due to the different positions of stimulus presentation in two experiments (right peripheral vs. central), in Experiment 1, the functional ROIs were identified within the left (i.e., contralateral) and right (i.e., ipsilateral) anatomical ROIs, respectively; and in Experiment 2, identified within the anatomical ROIs merged between hemispheres.

### fMRI analyses: MVPA

For Experiment 1, to examine how well the orientation of the motion path (left vs. right) of the perceived/imagined Gabor pattern could be discriminated by the multivoxel response patterns in different brain regions, we performed an ROI-based multivariate pattern analysis (MVPA), separately for each participant, using a multi-class support vector machine (SVM) classifier built-in in MATLAB R2018b (‘fitceoc’ and ‘predict’ functions). The SVM classifier was trained and tested on data within every condition, using a leave-one-run-out cross-validation procedure (i.e., patterns from all-but-one runs served as the training set, whereas patterns from the remaining run constituted the test set). In each condition, the decoding accuracies were obtained by calculating the proportion of trials in which the classifier correctly estimated the path orientation among all trials of the condition. Besides decoding of orientations within each condition, we also performed temporal generalization and cross-condition decoding analyses, using a similar leave-one-run-out procedure. Specifically, in temporal generalization analysis, for each condition, the classifier was trained on data from each time point, and tested on data from every time point. In cross-condition decoding analysis, the classifier was trained on data from each condition, and tested on data from all three conditions. The procedure of all decoding analyses described above was performed for each time point and ROI separately. Trials with opposite responses (i.e., responding “left” on right trials, or vice versa) were excluded from the decoding analyses. The remaining trials were further balanced between left and right trials within each condition to avoid overfitting in decoding. Overall, for each condition, 2.1 ± 1.5 trials were excluded after this procedure.

### fMRI analyses: IEM

For Experiment 2, to reconstruct the neural representation of orientation in each ROI, the inverted encoding model (IEM) analyses were implemented in MATLAB using custom codes (Brouwer and Heeger, 2009, 2011; Ester et al., 2015; Yu and Shim, 2017; Yu and Postle, 2021). The IEM assumes that the activation of each voxel could be characterized by the linear combination of a set of channel responses expressed by idealized tuning functions. Here, nine orientation channels equally spaced between 1-180° were used and the response of each channel was modelled as a sinusoid raised to the eighth power as follows:

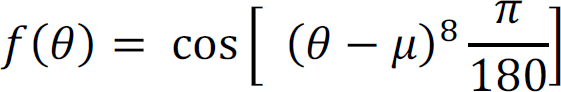

where θ is degrees in orientation space and μ is the channel center.

Independent training data B_1_ (*m* voxels × *n_1_* trials) were combined with the stimulus orientation channel profiles C_1_ (*k* channels × *n_1_* trials) to estimate an encoding model consisting of a weight matrix W (*m* voxels × *k* channels) which quantified the approximate contribution of each channel to the observed BOLD response in an ROI (i.e., B_1_). The relationship between B_1_, C_1_ and W can be described as follows:

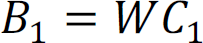

The weight matrix was computed through least-square linear regression as follows:

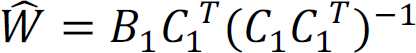

Following that, the encoding model was inverted such that it became a decoder that was used to estimate the channel responses C_2_ (*k* channels × *n_2_* trials) for the testing data B_2_ (*m* voxels × *n_2_* trials):

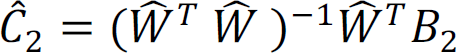

To make the estimated channel responses fully cover the stimulus space which comprised all integers ranging from 1° to 180°, the above steps were repeated 20 times and the centers of nine tuning functions were shifted by 1° on each iteration.

Finally, all estimated channel responses were circularly shifted to a common center (0° on the x axis representing the sample orientation of each trial) and averaged across trials, resulting in a single reconstruction for visualization and subsequent statistical comparisons. To quantify the strength of IEM reconstructions, the averaged channel responses on both sides of the common center were collapsed over and averaged, and the slope of each collapsed reconstruction was then computed through linear regression (Foster et al., 2017; Yu and Postle, 2021). A larger positive slope would indicate stronger positive representation.

The IEM analyses reconstructed population-level representations of orientations and allowed us to compare the orientation representational strength between different conditions within each ROI at any given time point. For a fair comparison between conditions to be made, we focused on the results of two IEM analyses where the IEM decoded testing data from different conditions with a common inverted encoder (Sprague et al., 2018): (1) a mixed IEM trained on data from all conditions (perception, imagery and illusion) and tested on data from each condition of the delayed-recall task; (2) an independent IEM trained on data averaged over 4.5-6 sec (4-5 th TRs; to avoid confusion, all the time points in the main text and methods referred to the onset of the corresponding TRs) of the perception-mapping task and tested on data from each condition of the delayed-recall task. Notably, the mixed IEM were trained and tested with a leave-one-run-out cross-validation procedure and the perception-mapping IEM was not. In addition, we also performed temporal generalization and cross-condition IEM analyses, following a similar procedure as those in Experiment 1. The procedure of all IEM analyses described above was performed for each time point and ROI separately.

### Statistical Procedures for multivariate analyses

For both experiments, a bootstrapping procedure was employed to evaluate the statistical significance of the accuracy/slope and also the statistical differences between accuracies/slopes. For Experiment 1, for each combination of factors (ROI, condition or TR), by randomly sampling with replacement from the pool of all 17 participants’ decoding results repeatedly, we generated 10000 simulated samples, each consisting of 17 decoding accuracies, and calculated the mean accuracy of each sample. The one-tailed p value of the decoding accuracy was computed to reflect the probability of obtaining a below-chance (< 0.5) accuracy among the 10,000 accuracies. For Experiment 2, for each combination of factors (ROI, condition, or TR), a bootstrapping procedure as above was performed with the IEM reconstructions, resulting in 10000 averaged IEM orientation reconstructions. Following that, the slope of each averaged reconstruction was computed and the proportion of negative slopes among the 10000 slopes was counted as the one-tailed *p* value of the slope. For time course of IEMs in each condition, the obtained *p* values denoting accuracy/slope significance were corrected for multiple comparisons using the FDR method across ROIs, conditions and time points. For temporal generalization analysis, the *p* values in the temporal generalization matrix were corrected using a cluster-based permutation method.

To better illustrate the accuracy/slope difference between conditions within different ROIs, we defined an early and a late epoch out of the sample and delay periods: for each ROI, the time points of the early epoch were chosen around the peak time points of the stimulus-evoked BOLD activity (Experiment 1: 4.5-6 s [4-5 th TRs] for EVC and 4.5-7.5 s [4-6 th TRs] for IPS; Experiment 2: 4.5-6 s [4-5 th TRs] for EVC and 6-7.5 s [5-6 th TRs] for IPS), whereas the time points of the late epoch were chosen as the last two TRs towards the end of the delay period (9-10.5 s [7-8 th TRs] for EVC and IPS in Experiments 1 and 2) during which cleaner memory-related signal was collected. For Figure 11, a common early epoch (4-6 th TRs) was used to allow direct comparisons between ROIs.

Within each epoch, we performed a two-way (ROIs × conditions) repeated measures ANOVA to test the differences between ROIs and conditions. For pairwise comparisons between conditions, we again employed a bootstrapping procedure. For instance, to test whether there were differences in accuracies between two conditions within a specific epoch and ROI, 17 pairs of accuracy were randomly sampled with replacement from the pool of 17 participants’ accuracies of these two conditions, and the difference between each pair was calculated and averaged. We repeated this procedure 10000 times, obtaining 10000 accuracy differences, and counted the proportion of accuracy differences with opposite signs among the 10,000 accuracy differences as the one-tailed *p* value of the accuracy difference between these two conditions. The significance of the slope difference between two IEM reconstructions was obtained using the same method described above. Within each epoch, *p* values were corrected for multiple comparisons using FDR across ROIs and comparisons.

### Univariate Analyses

To characterize univariate BOLD time course of each ROI during the delayed-recall task, we calculated the signal change in BOLD activity relative to baseline for each time point, where baseline was defined as the BOLD activity of the first TR of each trial. For each condition, the BOLD signal change was averaged across all voxels in each ROI and across trials within the condition. To test whether there was significant BOLD activity against baseline, one-tailed, one-sample *t* tests against 0 were used and the obtained *p* values were corrected across conditions and time points using the FDR method. To examine whether there were significant differences in BOLD activity between ROIs and conditions, we performed a two-way (ROIs × conditions) repeated measures ANOVA with each epoch. One-tailed, paired *t* tests were used and FDR corrected for subsequent pairwise comparisons between conditions.

### Eye-Tracking Data Analyses

Eye-tracking data were preprocessed using methods provided in (Nystrom and Holmqvist, 2010) and custom codes. Data were first filtered using a Savitzky-Golay (SG) FIR smoothing filter. Eye blinks and other artifacts were removed using a velocity-based algorithm and acceleration criteria. Data were then baseline corrected using average data from -200 to 0 ms prior to sample onset, and averaged within a 100-ms time window to reduced computational load. Because eye position data are only two-dimensional (x- and y-coordinates) and IEM thus does not apply, we instead used SVMs to decode orientations from eye position data. To be specific, we divided all trials into four bins based on their orientations (22.5-67.5°, 67.5-112.5°, 112.5-157.5°, 157.5-22.5°), and performed SVM classification between bins that were 90° apart. Decoding accuracy were then averaged across classifiers. This step was performed for each condition and each time point separately, and decoding results were further averaged across time points of sample and delay periods. Differences between eye position decoding performance in different conditions were evaluated using a one-way repeated measures ANOVA.

## Results

### Experiment 1

#### Behavior performance was comparable between perception, imagery, and illusion-based working memory in Experiment 1

Participants performed a delayed-recall task inside the MRI scanner, during which they memorized a physical, imagined, or illusory orientation of the motion path of a Gabor pattern over a prolonged delay as precisely as possible (Figure 1A). We referred these three conditions to perception-based, imagery-based, and illusion-based delayed-recall, respectively (perception, imagery and illusion in short, respectively). On perception trials, memoranda were Gabor patterns moving either along a leftward or rightward path in the right visual periphery. On imagery trials, no sample stimulus was presented, and participants had to imagine a leftward or rightward moving Gabor with the same orientations as those in the perception condition, based on a symbolic cue. On illusion trials, memoranda were Gabor patterns moving vertically, with an internal leftward or rightward drift. This type of double-drift stimulus would create a strong illusion on the perceived path of the motion, such that the perceived motion path was strongly deviated from its vertical physical path and tilted towards its internal drift direction (Liu et al., 2019). Prior to the main task, participants were measured on the size of their double-drift illusion (38.36° ± 5.71°). This measured illusion size was taken as the orientation of the motion path of the perception trials in the main task. In other words, the orientation of the motion path on the three conditions were designed to be matched within each participant, which we called the anchor orientation. In all conditions, participants reported their perceived or imagined orientation on an orientation wheel. Prior to the main experiment, participants filled out a revised version of the vividness of visual imagery questionnaire (VVIQ) (Marks, 1973) as an evaluation of their general imagery ability (1-5 points of rating: 1 – “no image at all”; 5 – “as vivid as normal vision”), with an average VVIQ score of 4.20 ± 0.55. Overall, participants’ performance was highly comparable across conditions, irrespective of the type of the sample stimulus (Table 1): The mean recall error, defined as the angular difference between reported and anchor orientations, did not significantly differ between conditions (*F*(2, 26) = 2.35, *p* = 0.116; one-way repeated measures ANOVA), neither did the standard deviation (SD) of the reported orientations (*F*(2, 26) = 1.46, *p* = 0.251; Figure 2A) nor the reaction time (RT) (*F*(2, 26) = 1.26, *p* = 0.300).

**Table 1.** Descriptive statistics of Experiments 1 and 2.

|  | Experiment 1 |  |  | Experiment 2 |  |
| --- | --- | --- | --- | --- | --- |
|  | Recall error<br>(Mean° ± SD°) | SD of<br>reported<br>orientations<br>(°) | Reaction<br>time<br>(Mean ± SD<br>in sec) | Recall error<br>(Mean° ± SD°) | Reaction<br>time<br>(Mean ± SD<br>in sec) |
| *Perception* | 6.35 ± 1.98 | 6.93 | 2.45 ± 0.32 | 6.70 ± 1.34 | 2.69 ± 0.29 |
| *Imagery* | 7.36 ± 2.37 | 7.91 | 2.50 ± 0.29 | 6.93 ± 1.10 | 2.72 ± 0.28 |
| *Illusion* | 6.77 ± 1.69 | 7.28 | 2.46 ± 0.25 | 7.04 ± 1.77 | 2.68 ± 0.24 |

**Figure 2.**
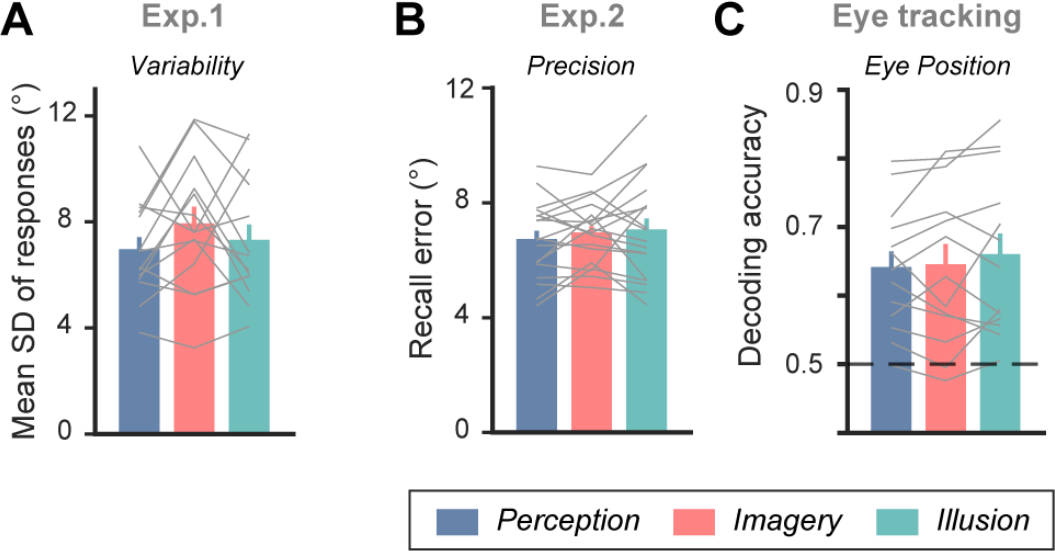
Behavioral results. A. Results of mean standard deviation of responses in each condition of Experiment 1. Colored bars indicate group mean (error bars denote ±1 SEM), gray lines indicate results from individual participants. B. Results of mean recall error in each condition of Experiment 2. Same conventions as A. C. Results of eye position decoding across sample and delay periods in the eye-tracking version of Experiment 2. Same conventions as A.

#### Orientation decoding in IPS was superior for imagery than perception in Experiment 1

To directly compare the representational strength of imagery and perception, here we focused on two pre-defined regions of interest (ROIs): low-level EVC (V1-V3) and interparietal sulcus (IPS) in higher-order parietal cortex. Because sample stimuli were presented in the right visual periphery, we focused all our primary analyses on contralateral ROIs, unless specified.

To investigate neural representations of stimulus-specific information in perception, imagery, and illusion, we performed Multivariate Pattern Analysis (MVPA) using Support Vector Machines (SVMs) to explore to what extent two path orientations (i.e., left vs. right) of the perceived/imagined stimuli could be discriminated on the basis of patterns of BOLD activity. Overall, significant and persistent above-chance classification performance was observed in all conditions and ROIs that began at slightly different time points (Figure 1B): In EVC, path orientations were decodable from sample onset on perception trials (*p*s < 0.040, tested using a bootstrapping procedure), earlier than that on imagery and illusion trials (from 3 sec onwards, *p*s < 0.034). By contrast, in IPS, decoding accuracy exceeded chance-level from 3 sec onwards on imagery trials (*p*s < 0.049), instead earlier than that on perception and illusion trials (from 4.5 sec onwards; *p*s < 0.032).

Given that EVC and IPS are highly reciprocally connected, any significant decoding result could be a result of spillover effects from downstream ROIs. Nevertheless, we hypothesized that if one brain region served as the source of decodable stimulus-specific signals in one specific condition, it should exhibit higher decoding performance in this condition compared to in other conditions. To test this hypothesis, we first defined an early and a late epoch out of the sample and delay period and, within each epoch, calculated the average decoding accuracy of each condition and ROI for further comparisons (see Methods for details). Our utilization of the delayed-recall task allowed for better dissociation between sample-evoked and memory-related signals. As such, results in the early epoch should mainly be accounted for by sample-driven signals, whereas results in the late epoch more likely reflected sustained, memory-related signals. During the early epoch (Figure 1C), a repeated-measures ANOVA demonstrated a significant interaction effect between conditions and ROIs (*F*(2, 32) = 48.09, *p* < 0.001). Pairwise comparisons further revealed a reversed pattern of accuracy differences between EVC and IPS as predicted: decoding accuracy was higher in perception compared to imagery in EVC (*p* < 0.001), whereas decoding accuracy was higher in imagery in IPS compared to perception (*p* = 0.004). Additionally, accuracy on illusion trials did not differ from imagery (*p* = 0.102), and was significantly lower than perception in EVC (*p* < 0.001); meanwhile, it did not differ from perception (*p* = 0.450), and was significantly lower than imagery in IPS (*p* = 0.002). Such a difference supported our hypothesis that illusion shared similar cognitive processes with imagery and perception at different stages of cortical processing. In comparison, during the late epoch, there were no significant differences between conditions (*p*s > 0.254; Figure 1D). To summarize, the fact that decoding performance for imagery was worse than perception in EVC, and better than perception and illusion in IPS, supported the notion that IPS might serve as the origin of stimulus-specific signals in imagery.

In Experiment 1, because only two path orientations were used, and sample stimuli between conditions differed dramatically in appearance, one might argue that the decodable signals were not specific to the path orientations, but instead reflected some more abstract signals such as semantic cues. To rule out this concern, we examined to what extent this imagery signal would be constrained retinotopically. We repeated the decoding analysis in the corresponding ipsilateral ROIs that received little bottom-up input signal, and found that, albeit classification performance was significantly above chance in all conditions (*p*s < 0.001), decoding accuracies did not differ between conditions (*p*s > 0.058, Figure 3). In other words, in ROIs outside the corresponding retinotopic areas of perceived/imagined location, the representational strength of path orientation during perception was similar to that during imagery. Note that our imagery cues were presented at fixation. As such, this retinotopic specificity of imagery provided further support that participants formed content-specific imagery at the corresponding retinotopic location following task instructions. Nevertheless, to fully address this concern as well as to test the generalizability of our findings, we conducted Experiment 2, in which participants perceived or imagined line orientations that continuously spanned the entire orientation space (1-180° in steps of 1°).

**Figure 3.**
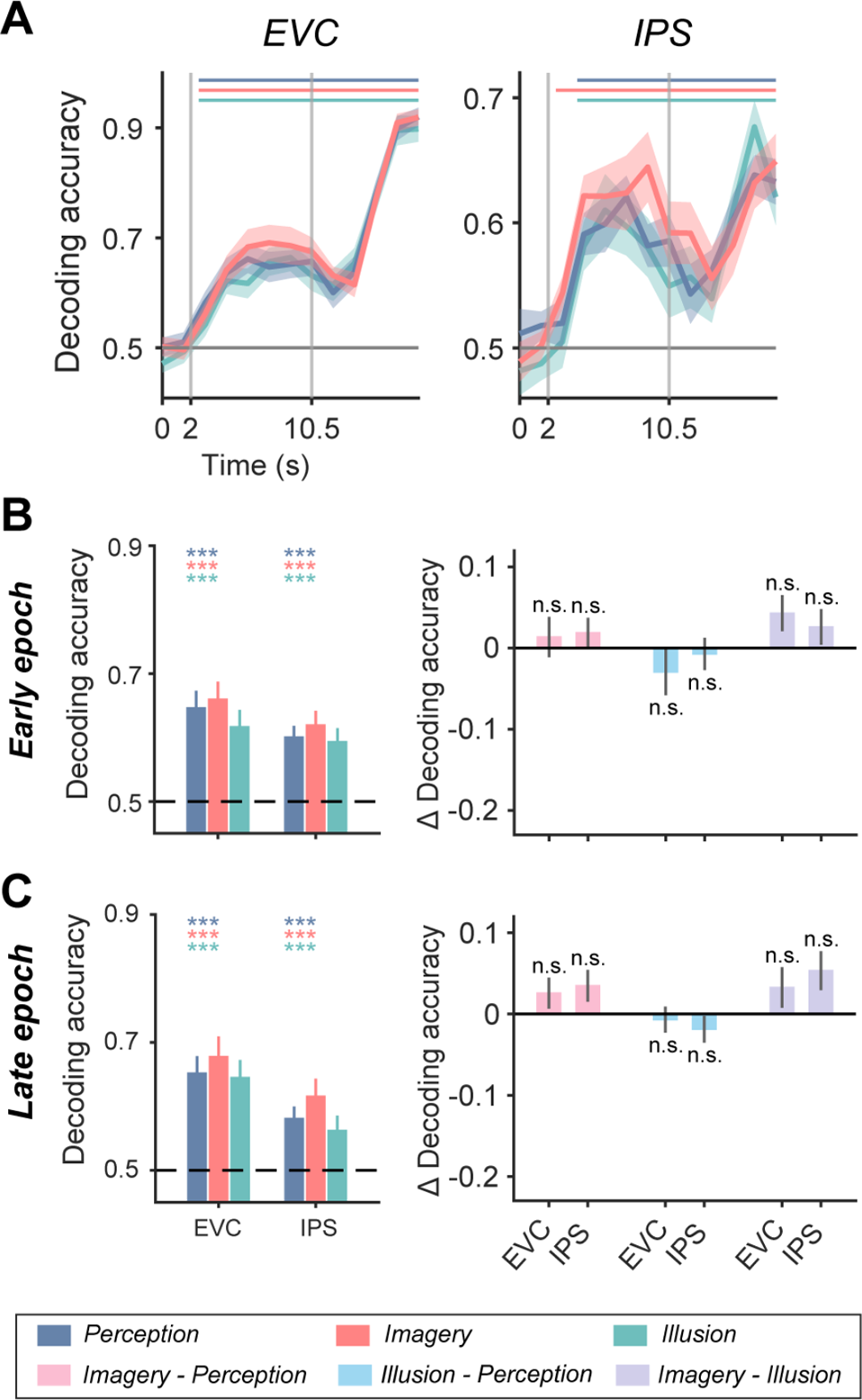
MVPA decoding of orientations in ipsilateral ROIs of Experiment 1. A. Time course of decoding accuracy in perception (blue), imagery (red), and illusion (green) conditions, in EVC and IPS of Experiment 1. ROIs were ipsilateral to the side of stimulus presentation. Colored lines on top denote significant time points of the corresponding condition. Vertical gray lines denote onset of delay (at 2 s) and of probe (at 10.5 s). Horizontal lines denote chance level of 0.5. Shaded areas denote error bars (±1 SEM). B. Left panel: average decoding accuracy in perception, imagery, and illusion conditions, in early epoch of EVC and IPS of Experiment 1. Horizontal dashed lines denote chance level of 0.5. Error bars denote ±1 SEM. Colored asterisks on top denote significance of the corresponding condition. Right panel: differences in decoding accuracy, between imagery and perception (pink), between illusion and perception (light blue), and between imagery and illusion (purple), in early epoch of EVC and IPS of Experiment 1. Error bars denote ±1 SEM. Asterisks denote significance of pairwise comparisons between conditions. n.s.: not significant, *: *p* < 0.05, **: *p* < 0.01, ***: *p* < 0.001. C. Same as B, but with results from late epoch.

### Experiment 2

#### Behavior performance was comparable between perception, imagery, and illusion-based working memory in Experiment 2

Participants performed an adapted version of delayed-recall task in Experiment 2: the trial structure was similar to that of Experiment 1, except that sample stimuli set was modified in order to trigger perceptual, imagined, and illusory experience of line orientations that covered the entire orientation space. The use of an entirely distinct stimuli set allowed us to directly examine whether the observed reverse pattern between EVC and IPS would hold with simple visual features, other than moving objects as in Experiment 1. Specifically, participants viewed a centrally-displayed sample stimulus for 1 sec (a bar with two end discs on perception trials; two distant discs on imagery trials; two distant discs with opposite openings on illusion trials) and then imagined or remembered the corresponding oriented bar over a 9.5-sec delay. After that, participants reported the orientation as precisely as possible (Figure 4A). Behavioral performance of Experiment 2 largely replicated that of Experiment 1: Participants reported an average VVIQ score of 3.96 ± 0.49, and no significant difference between conditions was observed in terms of recall error (*F*(2, 34) = 0.91, *p* = 0.413; Figure 2B) or reaction time (*F*(2, 34) = 1.95, *p* = 0.158; Table 1).

**Figure 4.**
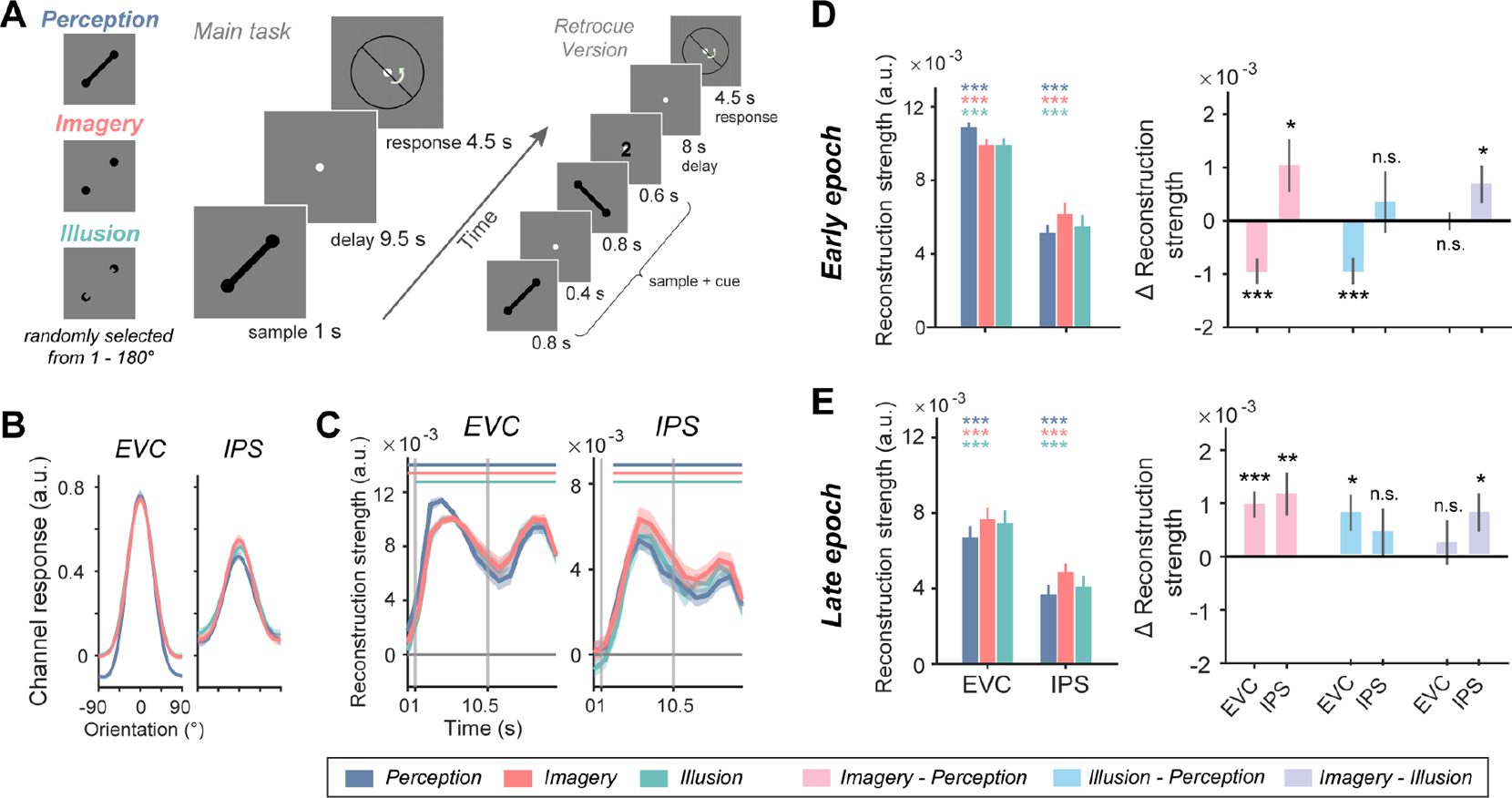
Task design and orientation representations in Experiment 2. A. In Experiment 2 (n = 18), the trial structure was similar to that of Experiment 1, except that sample stimuli sets were changed to static line orientations to cover the entire orientation space. Participants performed a delayed-recall task, during which they precisely memorized a physical, imagined, or illusory orientation over a prolonged delay. Each trial began with the presentation of one centrally-presented sample stimulus from one of the conditions. After a delay, participants rotated the needle presented at fixation to match the remembered orientation as precisely as possible. A subgroup of participants (n = 5) also performed a retrocue version of the task, in which two samples and a retrocue pointing to one of the samples were presented before the delay. B. Example orientation reconstructions using a mixed-IEM from selected time points with peak IEM reconstructions (4.5 s for EVC and 6 s for IPS), in perception, imagery, and illusion conditions, in EVC and IPS. X axis represents distance from sample orientations, with 0 representing the sample orientation of each trial. Y axis represents reconstructed orientation channel responses in arbitrary units. C. Time course of orientation reconstruction strength in Experiment 2. Colored lines on top denote significant time points of the corresponding condition. Vertical gray lines denote onset of delay (at 1 s) and of probe (at 10.5 s). Horizontal lines denote baseline of 0. Shaded areas denote error bars (±1 SEM). D. Left panel: average orientation reconstruction strength in perception, imagery, and illusion conditions, in early epoch of EVC and IPS of Experiment 2. Error bars denote ±1 SEM. Colored asterisks on top denote significance of the corresponding condition. Right panel: differences in orientation reconstruction strength, between imagery and perception (pink), between illusion and perception (light blue), and between imagery and illusion (purple), in early epoch of EVC and IPS of Experiment 2. Error bars denote ±1 SEM. Asterisks denote significance of pairwise comparisons between conditions. n.s.: not significant, *: *p* < 0.05, **: *p* < 0.01, ***: *p* < 0.001. E. Same as D, but with results from late epoch.

#### Orientation reconstructions in IPS were superior for imagery than perception in Experiment 2

Because all stimuli were centrally presented, in Experiment 2 we focused all neural analyses on bilateral ROIs. To characterize the neural representations of orientations on a continuous space, we implemented multivariate inverted encoding models (IEMs) to reconstruct population-level, orientation representations from voxel-wise brain activity within each ROI (Ester et al., 2015; Yu and Shim, 2017; Yu and Postle, 2021). We first trained a mixed IEM on data from all conditions and then tested on data from each condition at each time point, which allowed us to compare IEM reconstructions between task conditions in an unbiased way (Sprague et al., 2018). Figure 4B demonstrated example reconstructed orientation responses from selected time points in EVC and IPS, and the slope of each IEM reconstruction was computed to quantify the representational strength of orientation. For all conditions, the strength of orientation reconstructions arose markedly following stimulus presentation and remained above baseline over the delay in both EVC (from sample onset on imagery and perception trials, and from 1.5 sec on illusion trials, *p*s < 0.007) and IPS (from 3 sec; *p*s < 0.009; Figure 4C), indicating sustained representation of sample orientation in both visual and parietal cortex.

Next, similar to Experiment 1, an early and a late epoch were defined to better characterize the representational differences in visual and parietal cortex. During the early epoch, the interaction between ROI and condition was significant (*F*(2, 34) = 10.20, *p* < 0.001; Figure 4D). Subsequent pairwise comparisons confirmed the same reversed pattern in EVC and IPS between conditions as in Experiment 1, with EVC demonstrating stronger orientation representations on perception compared to imagery and illusion trials, and IPS demonstrating stronger orientation representations on imagery compared to perception and illusion trials (*p*s < 0.032). Moreover, orientation representations on illusion trials were more similar to those on imagery trials in EVC (*p*s > 0.307), and were more similar to those on perception trials in IPS (*p*s > 0.212) in both epochs, confirming that IPS differentiated between subjectively internal (i.e., imagery) and external (i.e., illusion and perception) experiences. Interestingly, during the late epoch (Figure 4E), the main effect of condition was significant (*F*(2, 32) = 6.09, *p* = 0.006), and orientation representations on imagery trials were stronger than those on perception trials in both EVC and IPS (*p*s < 0.003), and were stronger than those on illusion trials in IPS (*p* = 0.019). These results indicated that the enhanced imagery representations in IPS sustained into the late epoch, possibly implicating a sustained parietal signal in maintaining an internally-generated image in the task of Experiment 2.

### Enhanced orientation representations of imagery cannot be explained by univariate BOLD differences

Having identified enhanced orientation representations in IPS for imagery, at the multivariate level in both Experiments 1 and 2, we asked whether this result could be accounted for by differences in univariate BOLD activity. In Experiment 1, neither epoch demonstrated a significant interaction effect in BOLD activity (*F*s < 0.62, *p*s > 0.652), and sample presentation evoked stronger BOLD activity on perception and illusion compared to imagery trials in both ROIs (*t*s > 2.59, *p*s < 0.015; Figure 5A-C). In Experiment 2, both epochs demonstrated significant interaction effects (*F*s > 4.40, *p*s < 0.021). Nevertheless, There was no difference between three conditions ( *t*s < 1.85, *p*s > 0.227) but only a slightly higher response on imagery and illusion trials in IPS in early epoch (*t*s > 2.58, *p*s < 0.029; Figure 5D-F). In other words, multivariate and univariate analyses exhibited distinct patterns of condition differences, implicating that our decoding results were unlikely to be explained by univariate BOLD activation differences between conditions.

**Figure 5.**
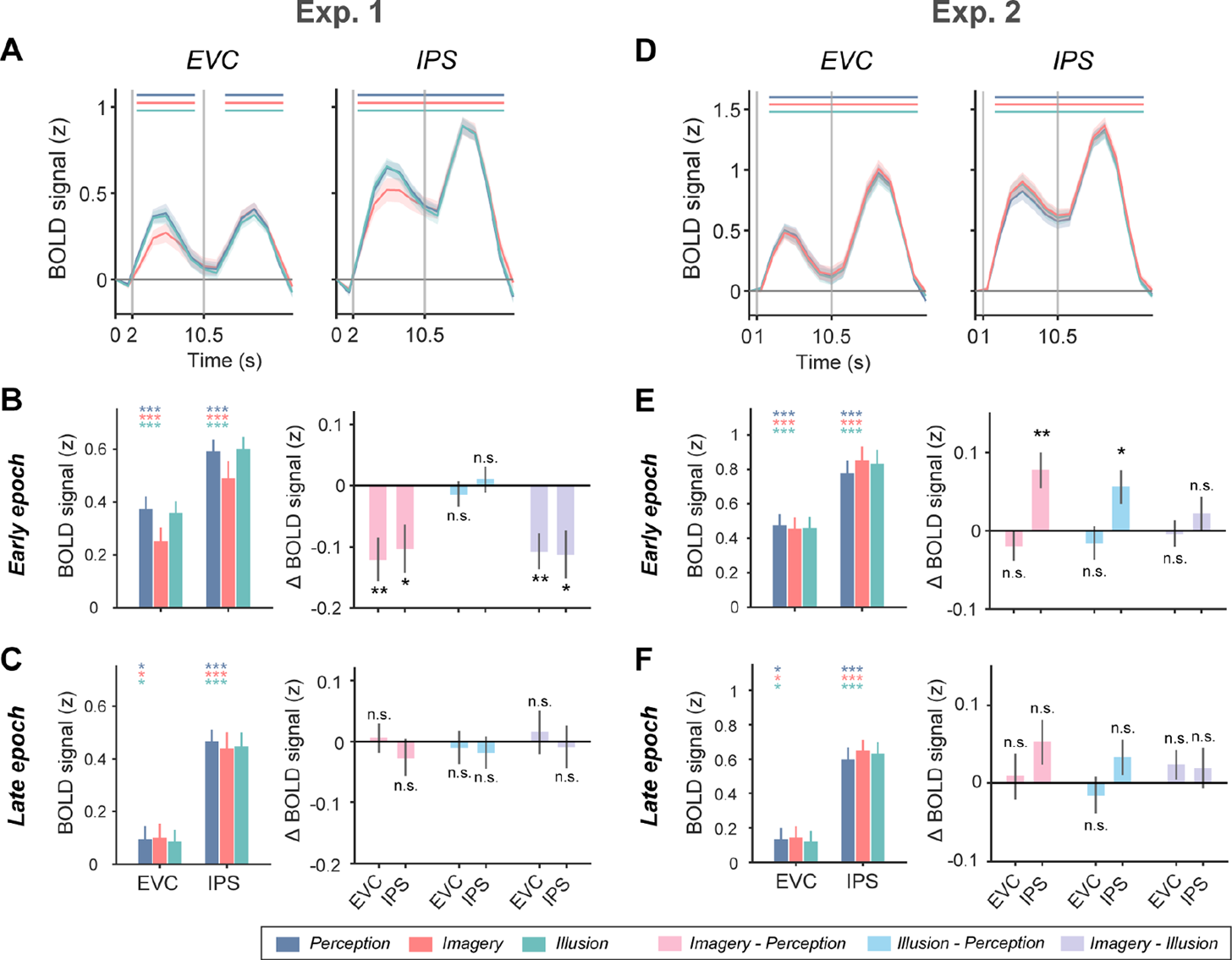
Univariate BOLD activity. A. Time course of BOLD signal change in perception (blue), imagery (red), and illusion (green) conditions, in EVC and IPS of Experiment 1. ROIs were contralateral to the side of stimulus presentation. Colored lines on top denote significant time points of the corresponding condition. Vertical gray lines denote onset of delay (at 2 s) and of probe (at 10.5 s). Horizontal lines denote baseline of 0. Shaded areas denote error bars (±1 SEM). B. Left panel: average BOLD signal change in perception, imagery, and illusion conditions, in early epoch of EVC and IPS of Experiment 1. Error bars denote ±1 SEM. Colored asterisks on top denote significance of the corresponding condition. Right panel: differences in BOLD signal change, between imagery and perception (pink), between illusion and perception (light blue), and between imagery and illusion (purple), in early epoch of EVC and IPS of Experiment 1. Error bars denote ±1 SEM. Asterisks denote significance of pairwise comparisons between conditions. n.s.: not significant, *: *p* < 0.05, **: *p* < 0.01, ***: *p* < 0.001. C. Same as B, but with results from late epoch of Experiment 1. D. Similar to A, but with results from Experiment 2. Vertical gray lines denote onset of delay (at 1 s) and of probe (at 10.5 s). E. Similar to B, but with results from Experiment 2. F. Similar to C, but with results from Experiment 2.

This was further validated by a series of control analyses: in Experiment 1, when we removed the mean BOLD activity from each condition and repeated the decoding analysis, similar patterns of results remained (Figure 6A-B). Likewise, controlling for the mean BOLD difference between left and right orientations within condition did not change the patterns of results either (Figure 6C-D), suggesting the decoding differences were exclusively linked to neural representations encoded in multivariate activation patterns. In Experiment 2, again when we removed the mean BOLD activity between conditions (Figure 7, the results largely remained (*p*s < 0.036, except for early epoch of IPS: imagery vs. perception, *p* = 0.092, imagery vs. illusion, *p* = 0.346). Hence, again the differences in representational strength across conditions could not simply be explained by differences in mean BOLD activity.

**Figure 6.**
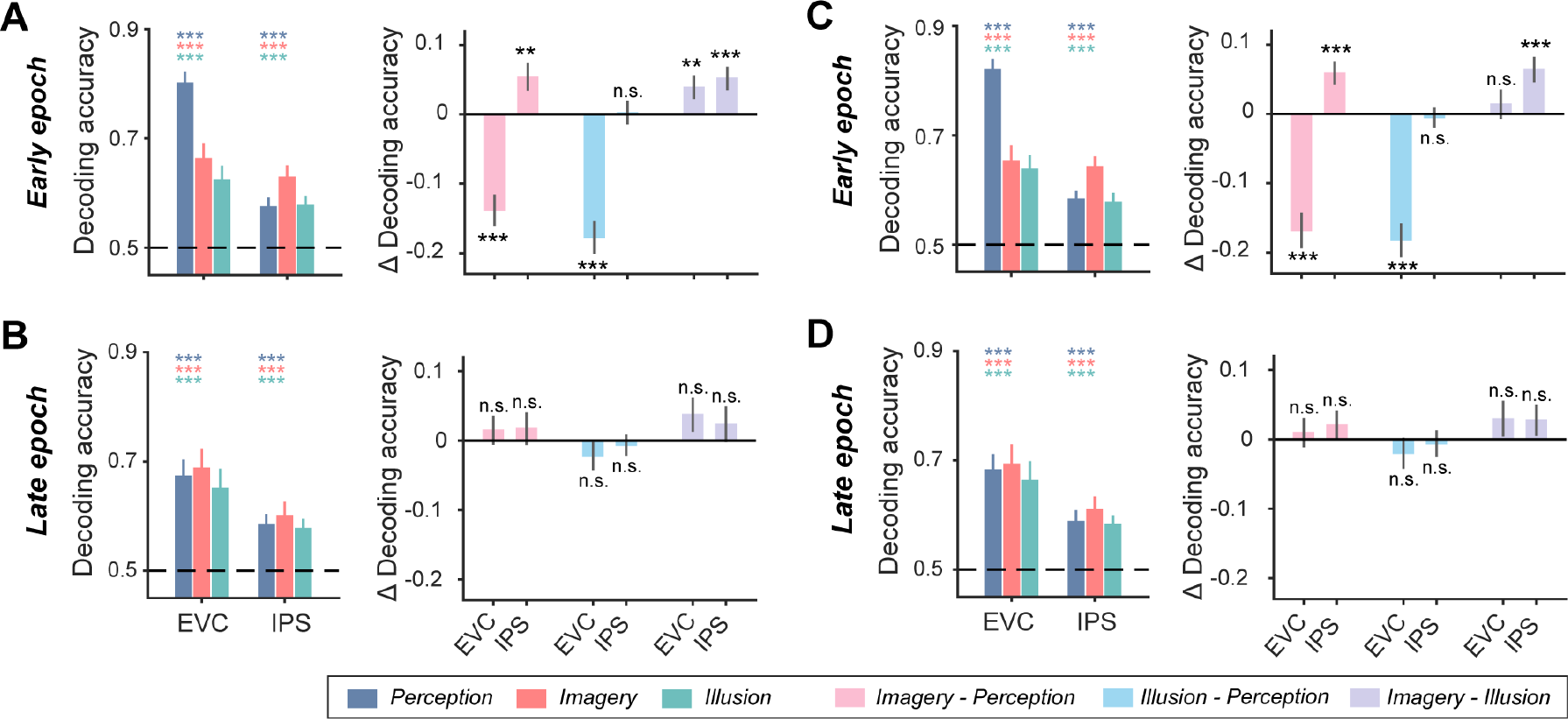
MVPA decoding results in Experiment 1 after controlling for differences in univariate BOLD activity. A. Left panel: average decoding accuracy in perception (blue), imagery (red), and illusion (green) conditions, in early epoch of EVC and IPS, after removing mean differences in BOLD activity between conditions. ROIs were contralateral to the side of stimulus presentation. Horizontal dashed lines denote chance level of 0.5. Error bars denote ±1 SEM. Colored asterisks on top denote significance of the corresponding condition. Right panel: differences in decoding accuracy, between imagery and perception (pink), between illusion and perception (light blue), and between imagery and illusion (purple), in early epoch of EVC and IPS of Experiment 1. Error bars denote ±1 SEM. Asterisks denote significance of pairwise comparisons between conditions. n.s.: not significant, *: *p* < 0.05, **: *p* < 0.01, ***: *p* < 0.001. B. Same as A, but with results from late epoch. C. Same as A, but with mean differences in BOLD activity between two path orientations within each condition removed. D. Same as B, but with mean differences in BOLD activity between two path orientations within each condition removed.

**Figure 7.**
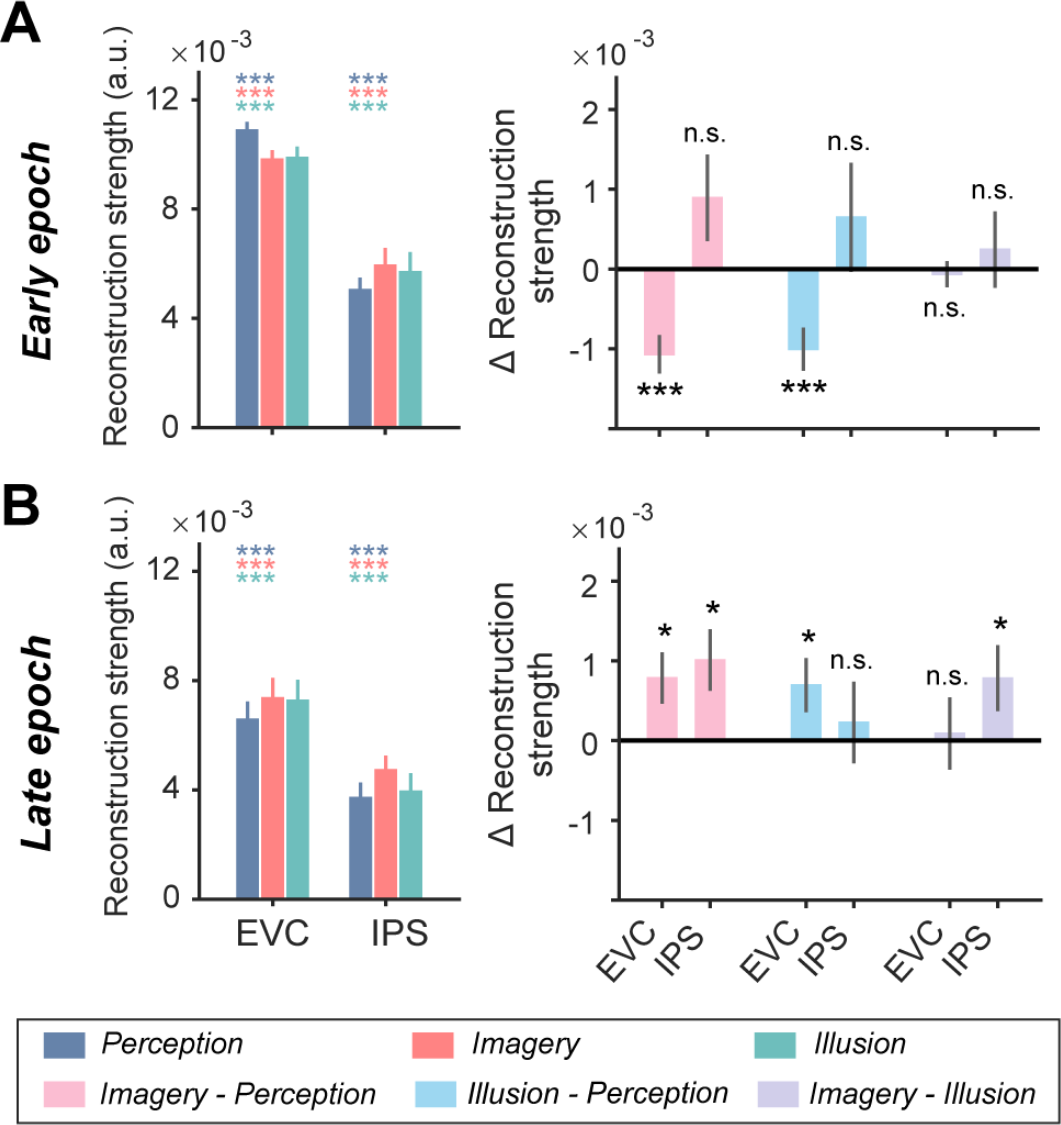
Orientation reconstructions in Experiment 2 after controlling for differences in univariate BOLD activity. A. Left panel: average orientation reconstruction strength in perception (blue), imagery (red), and illusion (green) conditions using a mixed-IEM, in early epoch of EVC and IPS, after removing mean differences in BOLD activity between conditions. Error bars denote ±1 SEM. Colored asterisks on top denote significance of the corresponding condition (against baseline of 0). Right panel: differences in orientation reconstruction strength, between imagery and perception (pink), between illusion and perception (light blue), and between imagery and illusion (purple), in early epoch of EVC and IPS. Error bars denote ±1 SEM. Asterisks denote significance of pairwise comparisons between conditions. n.s.: not significant, *: *p* < 0.05, **: *p* < 0.01, ***: *p* < 0.001. B. Same as A, but with results from late epoch.

### Enhanced orientation representations of imagery cannot be explained by differences in eye movement patterns

In Experiment 2, sample stimuli on imagery trials were two distant discs without a line presented at the fovea. This raised the possibility that participants might have had more eye movements towards the distant discs along the target orientation, thereby resulting in better neural representations of orientations on imagery trials. This was unlikely, given that better orientation representation was not observed in illusion, which should have also been influenced by eye movements, if existed. Nevertheless, to rule out this possibility, we conducted a behavioral control experiment during which we recruited a new group of volunteers (n = 13) to perform the same task as Experiment 2 with a shorter delay of 2.4 sec. Our results showed that decoding of orientations using eye position data revealed no significant differences between conditions (*F*(2, 24) = 1.06, *p* = 0.362; Figure 2C), suggesting our neural IEM results were unlikely to be contaminated by differences in stimulus-evoked eye movements.

### Enhanced orientation representations of imagery remained with a retrocue design

In both experiments, only one sample was presented and the sample differed in appearance between conditions, thereby resulting in inherently differential sample-evoked activity. Although we have demonstrated, through a series of univariate and multivariate analyses across ROIs, that our results were unlikely explained by physical differences in samples, we nevertheless performed a tentative study with a small group of participants (n = 5): the experimental procedure was identical to that in Experiment 2, except that now a retrocue paradigm was employed, such that participants viewed two sequentially presented samples at the beginning of each trial, and were retrocued on the to-be-maintained sample shortly after that. This retrocue design ensured that participants performed internal selection on the to-be-maintained orientation, rather than simply relying on stimulus difference between conditions. While this type of design significantly reduced stimulus-driven effects in EVC, the difference between imagery and perception remained in IPS, that is, orientation representation in imagery remained stronger than that in perception in IPS in both epochs (*p*s < 0.001, Figure 8). These results again confirmed the crucial role of IPS in generating stimulus-specific representations in imagery.

**Figure 8.**
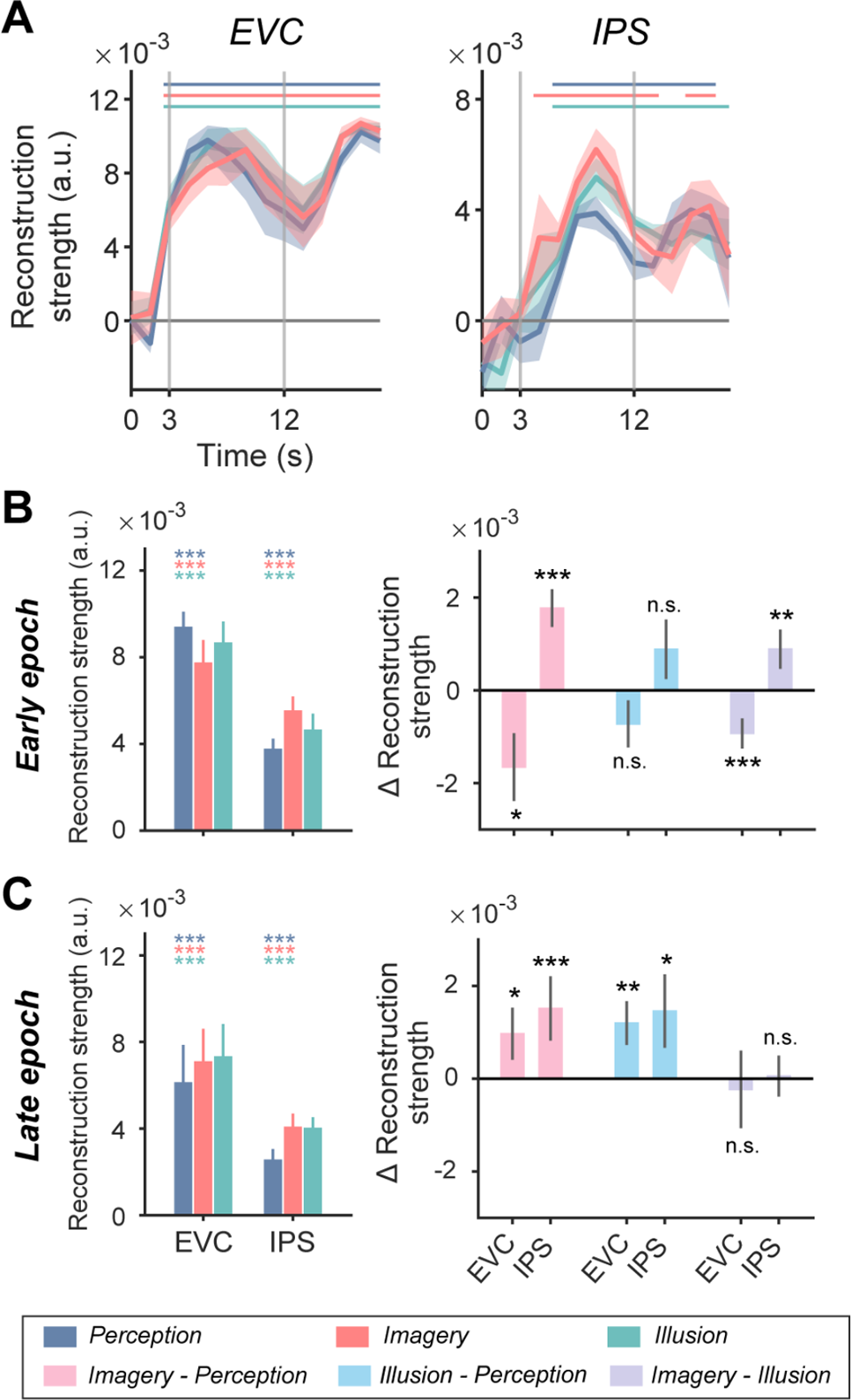
Orientation reconstructions in the retrocue version of Experiment 2. A. Time course of orientation reconstruction strength using a mixed-IEM in the retrocue version of Experiment 2. Colored lines on top denote significant time points of the corresponding condition. Vertical gray lines denote onset of delay (at 3 s) and of probe (at 12 s). Horizontal lines denote baseline of 0. Shaded areas denote error bars (±1 SEM). B. Left panel: average orientation reconstruction strength in perception, imagery, and illusion conditions, in early epoch of EVC and IPS of Experiment 2. Error bars denote ±1 SEM. Colored asterisks on top denote significance of the corresponding condition. Right panel: differences in orientation reconstruction strength, between imagery and perception (pink), between illusion and perception (light blue), and between imagery and illusion (purple), in early epoch of EVC and IPS of Experiment 2. Error bars denote ± 1 SEM. Asterisks denote significance of pairwise comparisons between conditions. n.s.: not significant, *: *p* < 0.05, **: *p* < 0.01, ***: *p* < 0.001. C. Same as B, but with results from late epoch.

### The representational format of orientation representations

Besides decoding performance of each time point, we further investigated the time-varying population dynamics in each condition, by examining the temporal generalization decoding patterns within each condition. In Experiment 1, we found that in EVC, neural codes in imagery and illusion were stable over time, while neural codes in perception were more dynamic, and underwent a clear transition from sample to delay period. In IPS, a similar transition was observed in imagery but not in perception or illusion (Figure 9A). These changes in population dynamics may constitute another neural signature that underlie the generation of imagery signals. In Experiment 2, however, neural codes in all three conditions remained stable over time, even in the perception condition (Figure 9C; this point will be elaborated in Discussion).

**Figure 9.**
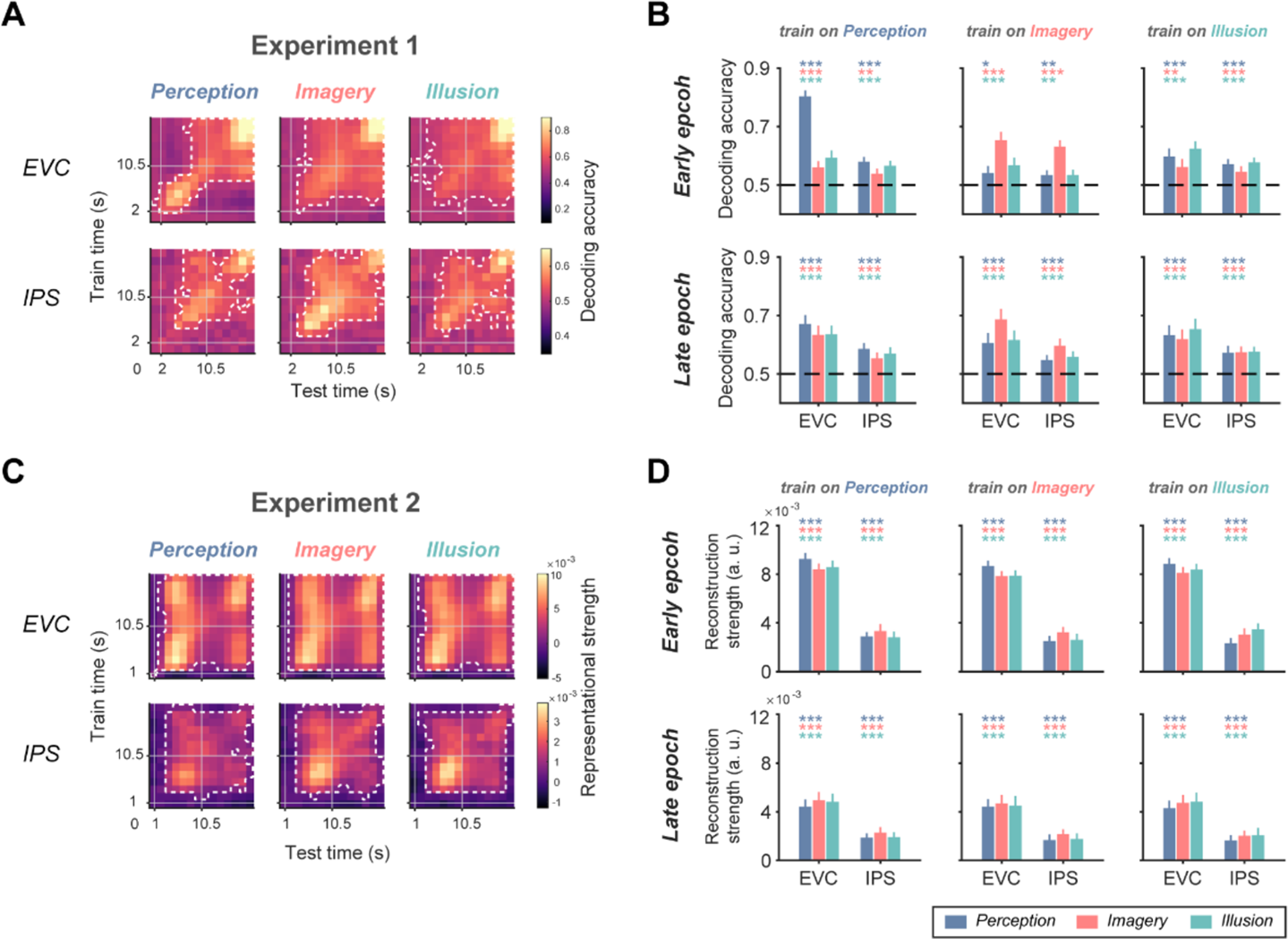
Temporal generalization and cross-condition results. A. Temporal generalization results in perception, imagery, and illusion conditions, in EVC and IPS of Experiment 1. ROIs were contralateral to the side of stimulus presentation. X and y axis denote test and train time, respectively. White dashed lines denote clusters with significant orientation representations after correction. B. Average decoding accuracy in cross-condition decoding of Experiment 1 when classifier was trained on one of the three conditions, and tested on all conditions, in EVC and IPS, for early epoch and late epoch separately. Colored asterisks on top denote significance of decoding performance of each condition, *: *p* < 0.05, **: *p* < 0.01, ***: *p* < 0.001. Horizontal dashed lines denote chance level of 0.5. Error bars denote ±1 SEM. C. Similar to A, but with results from Experiment 2. D. Similar to B, but with average orientation reconstruction strength in cross-condition IEM from Experiment 2.

In addition, to compare the representational format of orientations between conditions, we conducted a cross-condition decoding analysis, during which the classifier was trained on data from one condition and tested on the other two conditions separately for each epoch. We found that all three conditions were mutually generalizable in all ROIs during both epochs (*p*s < 0.018), in both Experiment 1 (Figure 9B) and Experiment 2 (Figure 9D), suggesting perception, imagery, and illusion shared common stimulus-specific representations in both EVC and higher-order IPS, although the relative representational strength differed between conditions within each ROI.

In Experiment 2, we collected data from an additional perception-mapping task on orientations, which allowed us to train a fixed model using the perception-mapping data and to make comparisons between conditions and epochs based on their neural similarities to perception. To be specific, we trained another IEM on data from the independent orientation perception-mapping task, and tested on data from each of the three conditions. Overall, results from this perception-mapping IEM were reminiscent of those from the mixed IEM: orientation representations were significant throughout the trial in EVC (*p*s < 0.001) and IPS (*p*s < 0.034), suggesting shared representations between internally-generated and externally-stimulated signals, and also between maintained and perceived signals (Figure 10A-B). Patterns of representational differences also largely replicated the main findings (*p*s < 0.049 except for early epoch of IPS: imagery vs. illusion, *p* = 0.219; Figure 10C-D). Note that stronger orientation representation on imagery trials was still observed in IPS when using the perception-mapping IEM. Together, these results further confirmed that IPS was more involved in imagery compared to perception.

**Figure 10.**
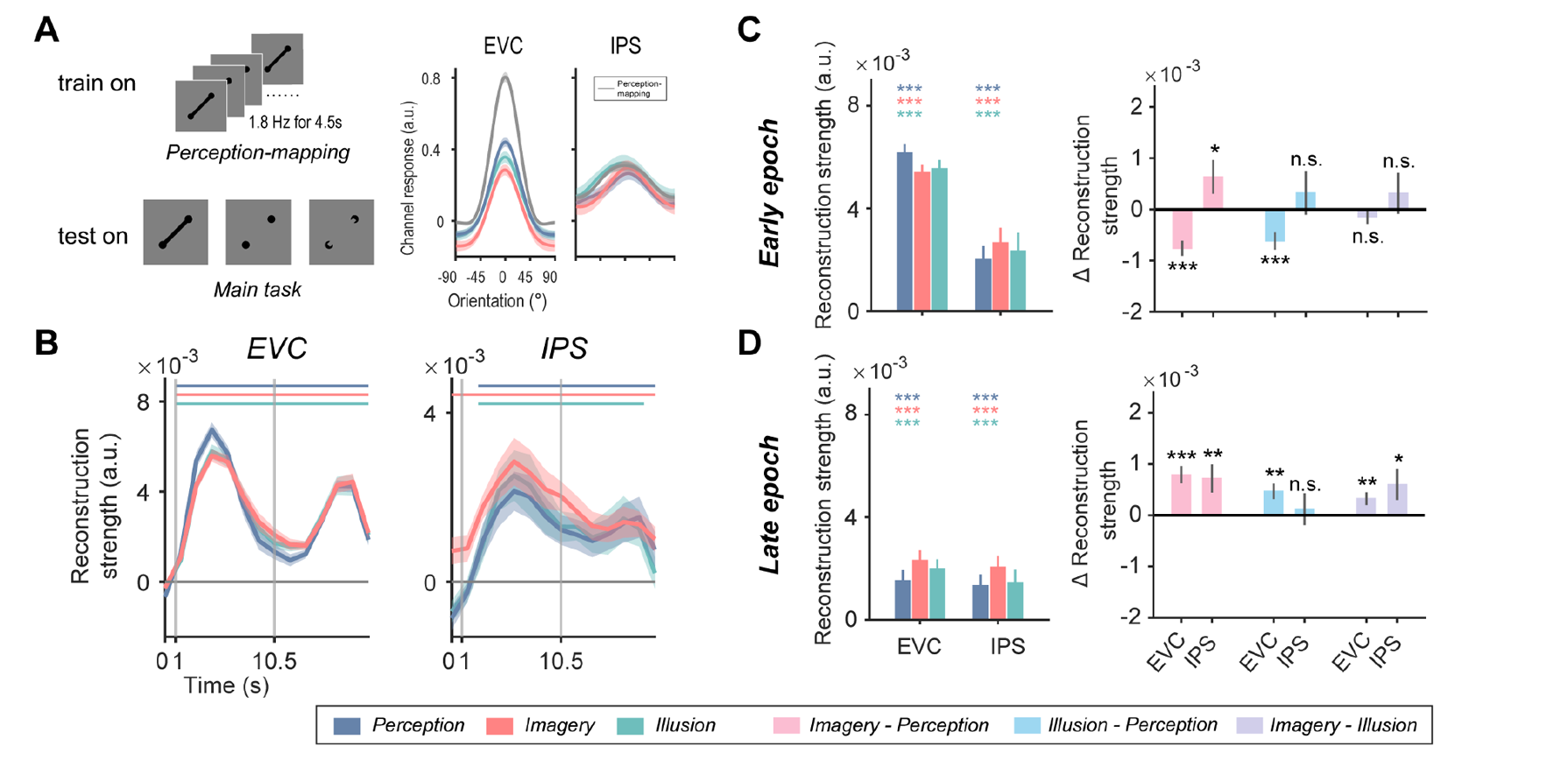
Orientation representations in Experiment 2 with a perception-mapping IEM. A. Example orientation reconstructions using a perception-mapping IEM from selected time points with peak IEM reconstructions (4.5 s for EVC and 6 s for IPS for main task, 4.5-6 s for perception mapping), in perception, imagery, and illusion conditions, in EVC and IPS. X axis represents distance from sample orientations, with 0 representing the sample orientation of each trial. Y axis represents reconstructed orientation channel responses in arbitrary units. B. Time course of orientation reconstruction strength in Experiment 2. Colored lines on top denote significant time points of the corresponding condition. Vertical gray lines denote onset of delay (at 1 s) and of probe (at 10.5 s). Horizontal lines denote baseline of 0. Shaded areas denote error bars (±1 SEM). C. Left panel: average orientation reconstruction strength in perception, imagery, and illusion conditions, in early epoch of EVC and IPS of Experiment 2. Error bars denote ±1 SEM. Colored asterisks on top denote significance of the corresponding condition. Right panel: differences in orientation reconstruction strength, between imagery and perception (pink), between illusion and perception (light blue), and between imagery and illusion (purple), in early epoch of EVC and IPS of Experiment 2. Error bars denote ±1 SEM. Asterisks denote significance of pairwise comparisons between conditions. n.s.: not significant, *: *p* < 0.05, **: *p* < 0.01, ***: *p* < 0.001. D. Same as C, but with results from late epoch.

### Emergence of imagery dominance along the posterior-to-anterior cortical hierarchy

We have provided converging evidence that IPS held more pronounced orientation representations during imagery, compared to perception and illusion. How did this imagery dominance emerge along the posterior-to-anterior cortical hierarchy, while this observation was absent in sensory-driven EVC? To address this question, we defined a set of functionally-activated, retinotopic ROIs from EVC to IPS, including V1, V2, V3, V3AB, IPS0, IPS1, IPS2, IPS3, IPS4, IPS5. In each retinotopic ROI, we calculated a difference index, between decoding accuracies of imagery and perception for Experiment 1, and between reconstruction strength of the two for Experiment 2. As depicted in Figure 11A, imagery dominance developed gradually along the EVC-IPS hierarchy in the early epoch in both experiments. Critically, an equilibrium point between imagery and perception was reached in V3AB, which lies in between EVC and IPS. In the late epoch of both experiments, enhanced orientation representations were observed during imagery in subregions of IPS, as well as in V3AB and/or EVC, suggesting an overall sustained enhancement of orientation representations in these regions during information maintenance of imagery.

**Figure 11.**
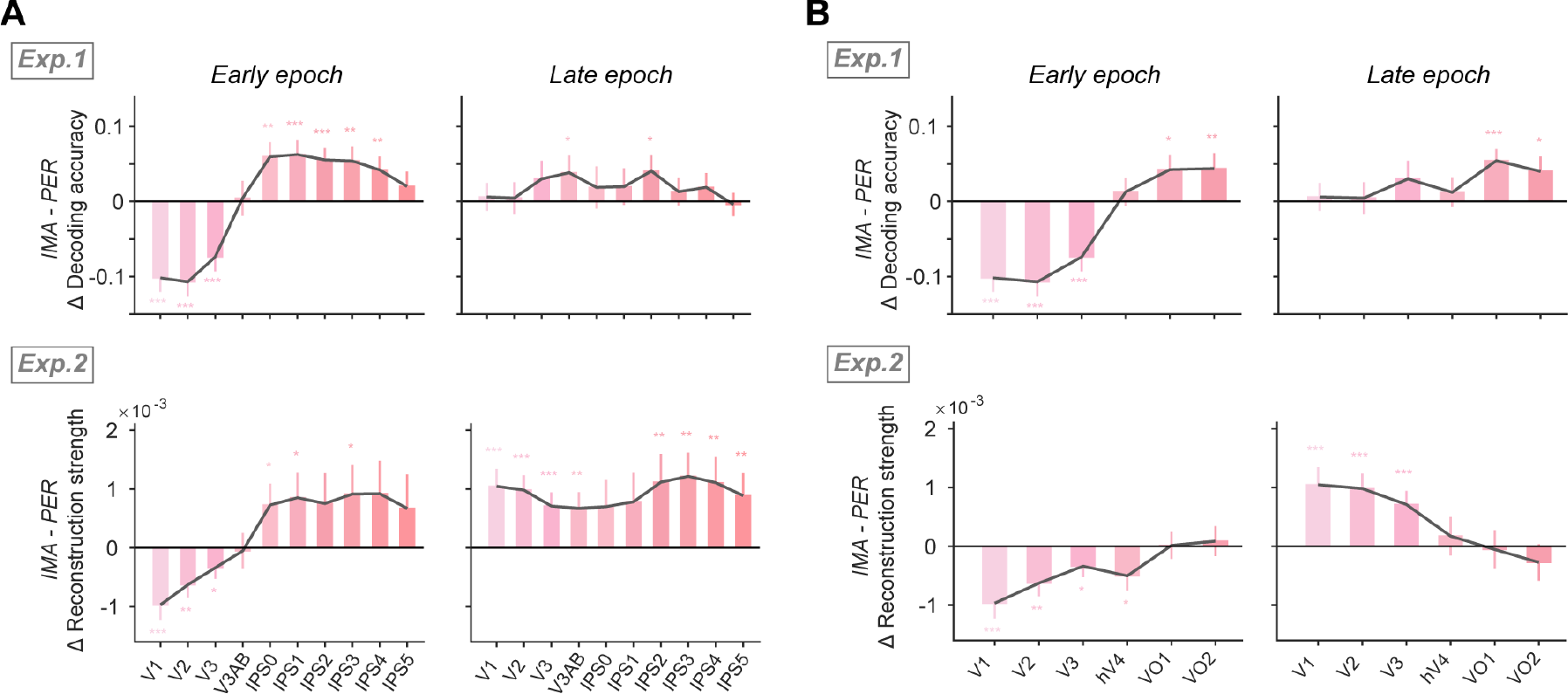
Relative representational strength changed along the retinotopic hierarchy. A. Differences in orientation representational strength between imagery and perception in retinotopic ROIs (V1-V3, V3AB, IPS0-5) along the EVC-IPS retinotopic hierarchy. Upper row: orientation decoding results from Experiment 1; Lower row: orientation reconstruction results from Experiment 2. Asterisks denote significant difference in the corresponding ROI, *: *p* < 0.05, **: *p* < 0.01, ***: *p* < 0.001. All *p* values remain uncorrected. Error bars denote ±1 SEM. B. Similar to A, but with results extended into the ventral visual stream (V1-V3, hV4, VO1, VO2).

### Only parietal cortex demonstrated a domain-general role in imagery

We have demonstrated superior orientation representations in imagery in parietal cortex across distinct stimuli sets (moving objects vs. visual feature), implicating a domain-general role of parietal cortex in imagery. How specific was this effect to parietal cortex? To address this question, we performed two additional analyses: first, we repeated the analysis in a series of ROIs along the ventral visual stream (hV4, VO1, VO2), and found that, enhanced representations for imagery were present in Experiment 1 where objects were used as stimuli, but were absent in Experiment 2 where only simple visual features were imagined (Figure 11B). Likewise, because moving objects were used in Experiment 1, we analyzed results from the motion-selective MT+ in both experiments. Similar to what we observed in the ventral visual stream, enhanced representations for imagery were found in MT+ in Experiment 1 (*p* < 0.001, Figure 12A) but not in Experiment 2 (*p* = 0.365, Figure 12B). These results together confirmed that object-selective and motion-selective visual areas exhibited a domain-specific role in imagery, as opposed to the domain-general role of parietal cortex.

**Figure 12.**
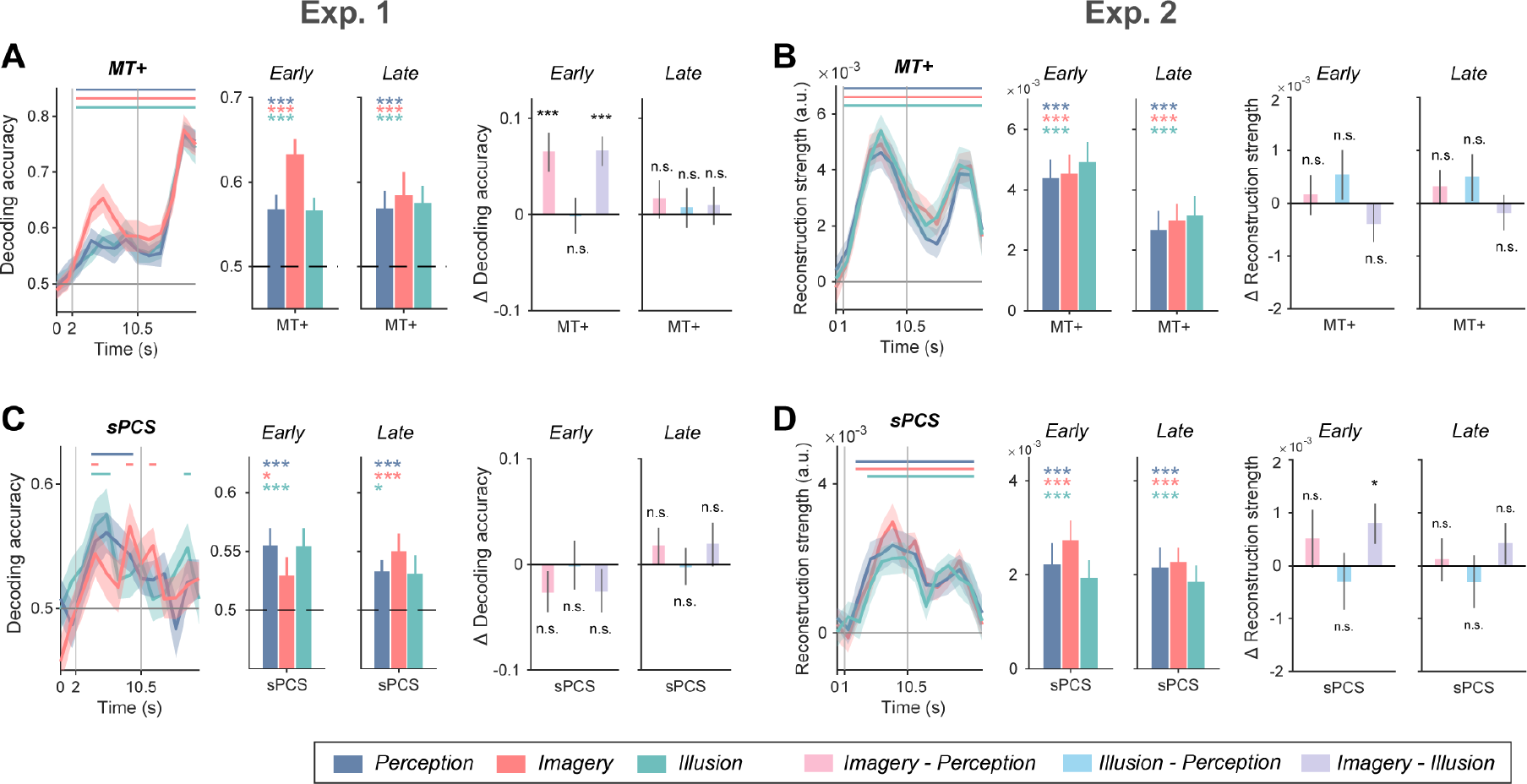
Orientation representations in MT+ and sPCS in Experiments 1 and 2. A. Left panel: time course of decoding accuracy in perception (blue), imagery (red), and illusion (green) conditions, in MT+ of Experiment 1. Colored lines on top denote significant time points of the corresponding condition. Vertical gray lines denote onset of delay (at 2 s) and of probe (at 10.5 s). Horizontal lines denote chance level of 0.5. Shaded areas denote error bars (±1 SEM). Middle panel: average decoding accuracy in perception, imagery, and illusion conditions. Horizontal dashed lines denote chance level of 0.5. Error bars denote ±1 SEM. Colored asterisks on top denote significance of the corresponding condition. Right panel: differences in decoding accuracy, between imagery and perception (pink), between illusion and perception (light blue), and between imagery and illusion (purple). Error bars denote ±1 SEM. Asterisks denote significance of pairwise comparisons between conditions. n.s.: not significant, *: *p* < 0.05, **: *p* < 0.01, ***: *p* < 0.001. B. Similar to A, but with results of orientation reconstruction strength using a mixed-IEM in MT+ of Experiment 2. (left) Vertical gray lines denote onset of delay (at 1 s) and of probe (at 10.5 s). Horizontal lines denote baseline of 0. C. Similar to A, but with results of sPCS in Experiment 1. D. Similar to C, but with results of sPCS in Experiment 2.

Second, we examined an additional ROI that lies more anterior to parietal cortex, i.e., the superior precentral sulcus (sPCS) in frontal cortex (Sprague and Serences, 2013; Yu and Shim, 2017), to see whether the observed patterns were widespread across multiple higher-order brain regions. We found that, sPCS failed to demonstrate an analogous pattern, in either Experiment 1 (*p*s > 0.148, Figure 12C) or Experiment 2 (*p*s > 0.249 except for imagery vs. illusion in early epoch: *p* = 0.040; Figure 12D). To summarize, the absence of differences in the strength of stimulus representations in brain regions with a higher cortical hierarchy than IPS, together with the positive evidence in IPS, implied a unique role of parietal cortex in forming stimulus-specific imagery.

## Discussion

In this study, we compared the neural processes underlying stimulus-specific representations in veridical perception, voluntary imagery and illusion. Across two experiments, population-level neural representations of stimulus-specific information in IPS discriminated between subjectively internal (imagery) and external (perception and illusion) experiences; whereas those in EVC discriminated between objectively internal (imagery and illusion) and external (perception) experiences. These results implicated IPS may serve as the source of content-specific imagery.

Previous neuroimaging work has demonstrated shared neural codes between imagery and perception in EVC (Albers et al., 2013; Ragni et al., 2020). However, it had remained less clear how stimulus-specific representations of imagery and perception differed from each other. Here we demonstrated superior stimulus-specific representations of perception than of imagery in EVC, consistent with the role of EVC in processing bottom-up external signals; By contrast, stimulus-specific representations of imagery were superior to those of perception in IPS. With the same logic, we interpreted this difference as reflecting a potential role of IPS in generating stimulus-specific imagery signals. We further demonstrated that this imagery dominance gradually developed along a functional gradient from EVC to IPS, with V3AB as an equilibrium point in between. Moreover, these results cannot be explained by differences in univariate BOLD activations between conditions, nor by influences from eye movements. Additionally, using a separate perception-mapping task to reconstruct orientation representations, and redoing Experiment 2 using a retrocue design, both yielded qualitatively similar results. Lastly, object/motion-selective regions demonstrated a domain-specific role as opposed to IPS, and examinations on a more anterior frontal region, sPCS, failed to replicate findings in IPS. To summarize, through a series of rigorous control experiments and analyses, we validated our findings of a unique imagery dominance in IPS.

Top-down modulation during imagery from parietal cortex has been supported by diverse neuroimaging evidence: in which parietal cortex shows consistent activation in imagery (Ishai et al., 2000; Ganis et al., 2004), content-independent connectivity with occipitotemporal areas (Mechelli et al., 2004), and significant correlation between BOLD activity and behavioral performance (Ragni et al., 2020). Nevertheless, the exact information being relayed from parietal cortex remains unclear. In particular, it has remained unclear whether imagined contents are robustly represented in parietal cortex, and how imagery representations in parietal cortex differ from perception (Albers et al., 2013; Ragni et al., 2020). Here, leveraging multivariate classification and IEMs, we provided converging evidence that imagery yielded significantly enhanced stimulus-specific representations in IPS than perception. The observation of robust imagery representations in IPS is in line with recent working memory studies demonstrating robust representations of memory contents in IPS (Ester et al., 2015; Yu and Shim, 2017; Rademaker et al., 2019; Yu and Postle, 2021), and provides new insights into the understanding of the nature of such representations: first, we demonstrated that stimulus-specific representations in IPS were not epiphenomenal, but rather functionally relevant and varied in strength with regard to the internality of information; second, we showed that enhanced representations of imagery emerged in the early epoch of the trial. These findings together suggested that internally-generated information originated from IPS. Additionally, we demonstrated that stimulus-specific representations in IPS, regardless of its exact format, were fine-tuned enough to allow successful reconstructions from a large set of different line orientations.

Compared to previous work, the current study included a delay period in all conditions. The memory delay allowed better dissociation between stimulus-evoked and memory-related signals in time. By this design, we were able to localize the representational differences to the sample epoch in both experiments. Moreover, similarly enhanced representations of imagery extended into the late delay epoch, suggesting IPS is involved in both the generation and maintenance of imagery representations. Although this imagery dominance in maintenance was relatively weaker and was only present in subregions of IPS in Experiment 1, it should be noted that a lack of imagery dominance in the late epoch does not necessarily mean that IPS did not contribute to imagery maintenance. Rather, we interpret the prolonged and pronounced effect in Experiment 2 as reflecting sustained efforts of IPS in maintaining imagery, possibly due to increased task difficulty brought by the large number of orientations used. Another advantage of including a memory delay was it allowed examination on the time-varying population dynamics of each condition (Spaak et al., 2017). Intriguingly, we observed a stable neural code across sample and delay periods in imagery and illusion in both experiments and in perception in Experiment 2, but a highly dynamic neural code in perception in Experiment 1. Perhaps relatedly, in Experiment 2, imagery and perception shared neural representations even in IPS, which was not the case between working memory and perception in several previous studies (Rademaker et al., 2019; Yu and Shim, 2019). We speculate that this apparent difference could be due to stimuli differences: we used more simplified stimuli (oriented lines) rather than gratings as in previous studies. The simplified stimuli might facilitate neural similarities in representing imagined and perceived contents (Kwak and Curtis, 2022), thereby resulting in better cross-condition and cross-time generalization. When using more complex stimuli such as those in Experiment 1, we observed a clear transition in neural code from sample to delay period in perception but not in imagery or illusion. Given that sample stimuli differed greatly in appearance in Experiment 1, it could be that the neural code during sample period reflected pure sensory-driven signals, and assimilated into a common neural code with imagery and illusion later into delay. This account would be consistent with previous work demonstrating absence of cross-decoding between physical and illusory stimuli in EVC when no memory was required (Liu et al., 2019). This difference in temporal dynamics might reflect another important neural signature that distinguishes imagery from perception.

Illusion can be conceptualized as misperception or involuntary imagery (Pearson and Westbrook, 2015; Pearson, 2019) within different theoretical framework. On one hand, illusion causes similar subjective experience as veridical perception that “information being perceived externally;” on the other hand, both illusory and imagined experience arise when the information is not directly accessible externally, yet illusory experience lacks the subjective feeling of “information being generated internally.” As such, illusion could serve as an ideal tool to dissociate subjective and objective internality. We found that, in EVC, illusion acted in a more comparable manner to imagery in terms of representational strength, reflecting a major contribution of physical sensory inputs regardless of subjective experience. In IPS, by contrast, the representational strength of orientation during illusion resembled that during perception to a larger extent, reflecting a dissociation between subjectively external and internal experiences. We argued that this result was unlikely due to an overall reduction in the strength of orientation representations during illusion, because representational strength in illusion did not significantly differ from that in imagery in EVC, nor from that in perception in IPS. In Experiment 2, representational strength in illusion was even numerically (but not statistically) higher in magnitude than in perception in several analyses. Collectively, these findings provided further support for our hypothesis that IPS carries neural signals that discriminate between subjectively internal and external experiences, which could be the critical reason of why imagery and perception feel so different.

Our results are broadly in line with the reverse hierarchy hypothesis, which proposes that imagery involves a reverse cortical hierarchy as opposed to perception, wherein signals originate from high-level frontoparietal and visual cortex and backpropagate to lower-level visual areas (Hochstein and Ahissar, 2002; Pearson, 2019). In addition, our findings add to the reverse hierarchy theory in several aspects: first, most of previous evidence in support of the reverse hierarchy focused on representational differences between low-level and high-level visual cortex along the ventral visual stream (Lee et al., 2012; Horikawa and Kamitani, 2017; Dijkstra et al., 2018; Dijkstra et al., 2020); and critically, representational strength of imagery never exceeded that of perception within the same brain region in those studies. Here, by having participants imagined or perceived orientation information, we attempted to minimize decoding difficulties between brain regions, thereby providing the first evidence for a representational “flip” between imagery and perception in parietal cortex. Second, we also observed enhanced imagery representations in object-selective visual cortex and motion-selective MT+ (Kaas et al., 2010) when using complex moving Gabors in Experiment 1. By contrast, in Experiment 2 when using simple features, the object/motion-related hierarchies disappeared, while the EVC-and-IPS hierarchy preserved. These results suggested that the source region of domain-specific imagery could be distributed in multiple areas and located flexibly in the cortical hierarchy, depending on the type of imagery contents. On the contrary, parietal cortex is involved in not only imagery of spatial information (Sack et al., 2005; Winlove et al., 2018), but also imagery of non-spatial stimuli such as features (Yu and Postle, 2021), objects (Dijkstra et al., 2017; Ragni et al., 2020; Ragni et al., 2021) and pictures (Breedlove et al., 2020). Therefore, parietal cortex may play a more domain-general role in visual imagery of various types of information. Whether the imagery dominance in parietal cortex generalizes to more types of information remains to be further elucidated in future research.

## Acknowledgements

We would like to thank Dr. Patrick Cavanagh for valuable comments on the manuscript, and Yiheng Hu for sharing part of the analysis codes. This work was supported by the National Natural Science Foundation of China (32271089), Shanghai Pujiang Program (22PJ1414400), the Ministry of Science and Technology of China (STI2030-Major Projects 2021ZD0204202, 2021ZD0203700), CAS Project for Young Scientists in Basic Research (YSBR-071), and Shanghai Municipal Science and Technology Major Project (2018SHZDZX05).

## Notes

### Competing Interest Statement

The authors have declared no competing interest.

## References

Albers AM, Kok P, Toni I, Dijkerman HC, de Lange FP (2013) Shared representations for working memory and mental imagery in early visual cortex. Curr Biol 23:1427–1431.

Arsenovic A, Ischebeck A, Zaretskaya N (2022) Dissociation between Attention-Dependent and Spatially Specific Illusory Shape Responses within the Topographic Areas of the Posterior Parietal Cortex. J Neurosci 42:8125–8135.

Bergmann J, Morgan AT, Muckli L (2019).

Breedlove JL, St-Yves G, Olman CA, Naselaris T (2020) Generative Feedback Explains Distinct Brain Activity Codes for Seen and Mental Images. Current Biology 30:2211–2224.e2216.

Brouwer GJ, Heeger DJ (2009) Decoding and reconstructing color from responses in human visual cortex. J Neurosci 29:13992–14003.

Brouwer GJ, Heeger DJ (2011) Cross-orientation suppression in human visual cortex. J Neurophysiol 106:2108–2119.

Cox RW (1996) AFNI: software for analysis and visualization of functional magnetic resonance neuroimages. Comput Biomed Res 29:162–173.

Dijkstra N, Bosch SE, van Gerven MA (2017) Vividness of Visual Imagery Depends on the Neural Overlap with Perception in Visual Areas. J Neurosci 37:1367–1373.

Dijkstra N, Ambrogioni L, Vidaurre D, van Gerven M (2020) Neural dynamics of perceptual inference and its reversal during imagery. Elife 9.

Dijkstra N, Mostert P, Lange FP, Bosch S, van Gerven MA (2018) Differential temporal dynamics during visual imagery and perception. Elife 7.

Ester EF, Sprague TC, Serences JT (2015) Parietal and Frontal Cortex Encode Stimulus-Specific Mnemonic Representations during Visual Working Memory. Neuron 87:893–905.

Foster JJ, Bsales EM, Jaffe RJ, Awh E (2017) Alpha-Band Activity Reveals Spontaneous Representations of Spatial Position in Visual Working Memory. Curr Biol 27:3216–3223 e3216.

Ganis G, Thompson WL, Kosslyn SM (2004) Brain areas underlying visual mental imagery and visual perception: an fMRI study. Brain Res Cogn Brain Res 20:226–241.

Hochstein S, Ahissar M (2002) View from the top: hierarchies and reverse hierarchies in the visual system. Neuron 36:791–804.

Horikawa T, Kamitani Y (2017) Generic decoding of seen and imagined objects using hierarchical visual features. Nat Commun 8:15037.

Iamshchinina P, Kaiser D, Yakupov R, Haenelt D, Sciarra A, Mattern H, Luesebrink F, Duezel E, Speck O, Weiskopf N, Cichy RM (2021) Perceived and mentally rotated contents are differentially represented in cortical depth of V1. Commun Biol 4:1069.

Ishai A, Ungerleider LG, Haxby JV (2000) Distributed neural systems for the generation of visual images. Neuron 28:979–990.

Kaas A, Weigelt S, Roebroeck A, Kohler A, Muckli L (2010) Imagery of a moving object: the role of occipital cortex and human MT/V5+. Neuroimage 49:794–804.

Kay L, Keogh R, Andrillon T, Pearson J (2022) The pupillary light response as a physiological index of aphantasia, sensory and phenomenological imagery strength. Elife 11.

Keogh R, Bergmann J, Pearson J (2020) Cortical excitability controls the strength of mental imagery. Elife 9.

Koenig-Robert R, Pearson J (2021) Why do imagery and perception look and feel so different? Philos Trans R Soc Lond B Biol Sci 376:20190703.

Kwak Y, Curtis CE (2022) Unveiling the abstract format of mnemonic representations. Neuron 110:1822–1828 e1825.

Lee SH, Kravitz DJ, Baker CI (2012) Disentangling visual imagery and perception of real-world objects. Neuroimage 59:4064–4073.

Lee SH, Kravitz DJ, Baker CI (2013) Goal-dependent dissociation of visual and prefrontal cortices during working memory. Nat Neurosci 16:997–999.

Liu S, Yu Q, Tse PU, Cavanagh P (2019) Neural Correlates of the Conscious Perception of Visual Location Lie Outside Visual Cortex. Curr Biol 29:4036–4044 e4034.

Marks DF (1973) Visual imagery differences in the recall of pictures. Br J Psychol 64:17–24.

Mechelli A, Price CJ, Friston KJ, Ishai A (2004) Where bottom-up meets top-down: neuronal interactions during perception and imagery. Cereb Cortex 14:1256–1265.

Nystrom M, Holmqvist K (2010) An adaptive algorithm for fixation, saccade, and glissade detection in eyetracking data. Behav Res Methods 42:188–204.

Pearson J (2019) The human imagination: the cognitive neuroscience of visual mental imagery. Nat Rev Neurosci 20:624–634.

Pearson J, Westbrook F (2015) Phantom perception: voluntary and involuntary nonretinal vision. Trends Cogn Sci 19:278–284.

Rademaker RL, Chunharas C, Serences JT (2019) Coexisting representations of sensory and mnemonic information in human visual cortex. Nat Neurosci 22:1336–1344.

Ragni F, Lingnau A, Turella L (2021) Decoding category and familiarity information during visual imagery. Neuroimage 241:118428.

Ragni F, Tucciarelli R, Andersson P, Lingnau A (2020) Decoding stimulus identity in occipital, parietal and inferotemporal cortices during visual mental imagery. Cortex 127:371–387.

Reddy L, Tsuchiya N, Serre T (2010) Reading the mind’s eye: decoding category information during mental imagery. Neuroimage 50:818–825.

Roelfsema PR, de Lange FP (2016) Early Visual Cortex as a Multiscale Cognitive Blackboard. Annu Rev Vis Sci 2:131–151.

Sack AT, Camprodon JA, Pascual-Leone A, Goebel R (2005) The dynamics of interhemispheric compensatory processes in mental imagery. Science 308:702–704.

Spaak E, Watanabe K, Funahashi S, Stokes MG (2017) Stable and Dynamic Coding for Working Memory in Primate Prefrontal Cortex. J Neurosci 37:6503–6516.

Sprague TC, Serences JT (2013) Attention modulates spatial priority maps in the human occipital, parietal and frontal cortices. Nat Neurosci 16:1879–1887.

Sprague TC, Adam KCS, Foster JJ, Rahmati M, Sutterer DW, Vo VA (2018) Inverted Encoding Models Assay Population-Level Stimulus Representations, Not Single-Unit Neural Tuning. eNeuro 5.

Stokes M, Thompson R, Cusack R, Duncan J (2009) Top-Down Activation of Shape-Specific Population Codes in Visual Cortex during Mental Imagery. Journal of Neuroscience 29:1565–1572.

Wang L, Mruczek RE, Arcaro MJ, Kastner S (2015) Probabilistic Maps of Visual Topography in Human Cortex. Cereb Cortex 25:3911–3931.

Winlove CIP, Milton F, Ranson J, Fulford J, MacKisack M, Macpherson F, Zeman A (2018) The neural correlates of visual imagery: A co-ordinate-based meta-analysis. Cortex 105:4–25.

Yu Q, Shim WM (2017) Occipital, parietal, and frontal cortices selectively maintain task-relevant features of multi-feature objects in visual working memory. Neuroimage 157:97–107.

Yu Q, Shim WM (2019) Temporal-Order-Based Attentional Priority Modulates Mnemonic Representations in Parietal and Frontal Cortices. Cereb Cortex 29:3182–3192.

Yu Q, Postle BR (2021) The Neural Codes Underlying Internally Generated Representations in Visual Working Memory. J Cogn Neurosci:1–16.

